# PMAIP1-Mediated Glucose Metabolism and its Impact on the Tumor Microenvironment in Breast Cancer: Integration of Multi-Omics Analysis and Experimental Validation

**DOI:** 10.1101/2024.10.30.620466

**Authors:** Yidong Zhang, Xuedan Han, Qiyi Yu, Lufeng Zheng, Hua Xiao

## Abstract

**Background:** Glucose metabolism in breast cancer has a potential effect on tumor progression and is related to the immune microenvironment. Thus, this study aimed to develop a glucose metabolism– tumor microenvironment score to provide new perspectives on breast cancer treatment.

**Method:** Data were acquired from the Gene Expression Omnibus and UCSC Xena databases, and glucose-metabolism-related genes were acquired from the Gene Set Enrichment Analysis database. Genes with significant prognostic value were identified, and immune infiltration analysis was conducted, and a prognostic model was constructed based on the results of these analyses. The results were validated by in vitro experiments with MCF-7 and MCF-10A cell lines, including expression validation, functional experiments, and bulk sequencing. Single-cell analysis was also conducted to explore the role of specific cell clusters in breast cancer, and Bayes deconvolution was used to further investigate the associations between cell clusters and tumor phenotypes of breast cancer.

**Results:** Four significant prognostic genes (PMAIP1, PGK1, SIRT7, and SORBS1) were identified, and, through immune infiltration analysis, a combined prognostic model based on glucose metabolism and immune infiltration was established. The model was used to classify clinical subtypes of breast cancer, and PMAIP1 was identified as a potential critical gene related to glucose metabolism in breast cancer. Single-cell analysis and Bayes deconvolution jointly confirmed the protective role of the PMAIP1+ luminal cell cluster.

## 1 Introduction

Breast cancer is among the most prevalent malignant cancers and the leading cause of cancer-associated deaths in women worldwide [1–3]. It has been characterized as a solid tumor with high heterogeneity, and more than 20 subgroups, including luminal A, luminal B, HER2, and triple-negative breast cancer, have been identified [4, 5]. The diverse biological subtypes of breast cancer differ in morphology, clinical subtypes, treatment response, and disease behaviors, posing a substantial challenge in diagnosis and treatment [6, 7]. Despite achievements in prevention, early detection, and personalized therapy for breast cancer—for example, new therapeutic targets such as SERPINA5 [8]—the mortality rate of breast cancer patients remains high, owing to tumor metastasis, chemotherapy resistance, and disease recurrence [9–11]. Thus, more effective prognostic biomarkers and therapeutic strategies are urgently needed to improve the efficacy of breast cancer treatment.

Alterations in energy metabolism, especially glucose metabolism, are typical characteristics of tumors [12–14]. These changes, known as the “Warburg effect”, involve the abnormal activity of glycolysis process to produce lactate, even in oxygen-enriched environments [15]. As a consequence, excessive consumption of oxygen and accumulated lactate contribute to a highly hypoxic and acidic microenvironment. In breast cancer, cellular aerobic glycolysis has been reported to contribute to tumor progression and metastasis; therefore, the glucose metabolism process has potential to be a significant prognostic factor in breast cancer [16, 17]. However, the specific characteristics of glycolytic metabolism in breast cancer are not clear. In addition, glucose metabolism characteristics vary among different cell clusters of breast cancer, resulting in complex metabolic phenotypes.

The tumor microenvironment (TME) is complex and variable, comprising various types of cells including tumor-infiltrating lymphocytes, immune cells, and fibroblasts, as well as extracellular matrix. Previous studies have investigated the associations between glucose metabolism of tumor cells and immune cell function [18]. These associations indicate that the highly hypoxic and acidic microenvironment resulting from tumor glycolysis promotes immune escape and development of cancer [19, 20]. Thus, elucidation of the part played by modulation of glucose metabolism in the establishment of the TME immune landscape in breast cancer could lead to optimization of clinical immune therapies and the discovery of new treatment targets.

The rapid development of bioinformatics methods has provided a new perspective on glucose metabolism in breast cancer. In the present study, significant glucose-metabolism-related genes and critical immune cells in breast cancer were identified via gene expression profile analysis, and a combined prognostic model was established. PMAIP1 (phorbol-12-myristate-13-acetate-induced protein 1) was identified as the critical hub gene of breast cancer prognosis associated with glucose metabolism, and its regulation to tumor phenotypes were validated in a breast cancer cell line. Bulk sequencing (bulk-seq) further revealed variations in glucose metabolism at the cell line level. Moreover, single-cell analysis was used to explore cell clusters in breast cancer, and the associations between PMAIP1+ luminal cells and tumor phenotypes in breast cancer were further investigated via Bayes deconvolution. In future work, we aim to explore and validate the significant glucose-metabolism-related genes in the progression of breast cancer and expect to provide novel potential therapeutic targets for further exploration.

## 2 Materials and Methods

### 2.1 Data acquisition

We acquired The Cancer Genome Atlas Breast Invasive Carcinoma (TCGA-BRCA) dataset, which included RNA sequencing (RNA-seq) gene profiles and clinical data of 1,217 breast cancer patients, from the UCSC Xena database (https://xenabrowser.net/datapages/). The dataset was analyzed in transcripts per million reads. The expression matrix of TCGA-BRCA was first annotated using the GRCh38 human reference genome profile; then, protein-coding genes were selected for further downstream analysis. If multiple probes were annotated as one gene symbol, the maximum expression level detected by probes was identified as the expression level of the gene. Survival information and clinical characteristics, including overall survival time, survival status, age, gender, M stage, N stage, T stage, and tumor stage were extracted from the patient clinical information as indices for clinical subtype classification. Patients with missing values were excluded.

In addition, we downloaded RNA-seq microarray expression profiles (Illumina HT-12 v3 microarray) and clinical data for 1,963 breast tumor samples and 17 control samples from the Molecular Taxonomy of Breast Cancer International Consortium (METABRIC) [21] and obtained breast cancer 10X Genomics single-cell sequencing dataset GSE195861 and microarray validation dataset GSE42568 from the Gene Expression Omnibus [22, 23]. GSE195861 included 13 breast cancer samples, six lymph node metastasis samples, and one normal breast tissue sample from the GPL20795 platform. GSE42568 comprised 104 breast cancer biopsies and 17 normal breast biopsies from the GPL570 platform.

### 2.2 Establishment of glucose-related score (GLC-Score)

We acquired the “GOBP_GLUCOSE_METABOLIC_PROCESS” gene set from the Gene Set Enrichment Analysis database and extracted a total of 193 glucose-metabolism-related genes [24, 25]. Based on this glucose metabolism gene set, we screened differentially expressed genes (DEGs) between breast cancer and control samples using the TGCA-BRCA expression profiles with the “limma” package (v. 3.54.0), with a threshold of adjusted p<0.05 [26]. After identifying DEGs, we performed univariate Cox regression analysis to identify prognosis-related DEGs in breast cancer. Subsequently, least absolute shrinkage and selection operator (LASSO) was used to select hub prognostic genes. The optimal λ parameter was determined by ten-fold cross validation. To evaluate the prognostic value of the selected genes and establish the GLC-Score, we used the “bootstrap_multicox” method was applied, performing 1,000 bootstrap iterations. Coefficient was extracted from the bootstrap progression, and the standard deviation (SD) values of coefficients were calculated for each selected gene. A GLC-Score for each sample was determined as the sum of coefficients divided by SD values from the bootstrap analysis multiplied by gene expression: 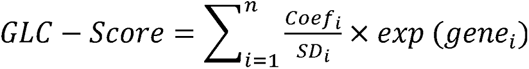. Parameter i represents the number of each selected genes, n represents the count of all selected genes. The acquired GLC-Scores were used to divide samples into GLC-High and GLC-Low groups according to the median value. Kaplan–Meier survival analysis was then conducted to determine the survival rates of breast cancer patients in the two groups.

### 2.3 Establishment of TME-score and combined prognostic analysis

A TME-Score was established based on immune infiltration in breast cancer. The fractions of 22 types of immune cell in each sample were estimated using the CIBERSORT method (R Script v. 1.03, obtained from the CIBERSORT website; https://cibersort.stanford.edu/) [27]. Survival analysis was then conducted to identify immune cells with significant protective effects on patient survival. The optimized cutoff point for stratification of samples was determined using the “surv_cutpoint” function. Subsequently, “bootstrap_multicox” analyses were performed to evaluate the risk posed by these immune cells to breast cancer patients. Coefficient was extracted from the bootstrap progression, and the standard deviation (SD) values of coefficients were calculated for each selected immune cell type. The corresponding TME-Score was determined as the sum of the coefficients divided by the SD values from the bootstrap analysis, multiplied by the cell proportion: 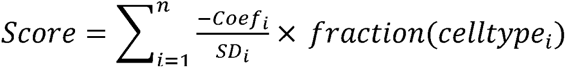. Parameter i represents the number of each selected immune cell type, n represents the count of all selected immune cell types. Kaplan–Meier survival analyses were performed to evaluate the survival of breast cancer patients, and patients were divided into TME-High and TME-Low groups based on the median value of the TME-Score.

Having established the GLC-Score and TME-Score, we constructed a combined prognostic model. Samples were divided into four groups, integrating the GLC-Score and TME-Score groups, as follows: GLC-High+TME-High, GLC-High+TME-Low, GLC-Low+TME-High, and GLC-Low+TME-Low. The survival of patients in these four groups was then analyzed, and receiver operating characteristic curves were used to measure the predictive ability of the prognostic model for 3-, 5-, and 7-year survival.

### 2.4 Weighted correlation network analysis (WGCNA) and functional enrichment

We used the WGCNA package (v. 1.72-1) to create a scale-free mRNA co-expression network in TCGA-BRCA dataset on the basis of the four groups described above [28]. First, we calculated the Pearson correlation coefficients of gene pairs to construct a similarity matrix, which we then transformed into an adjacency matrix. Using the criterion of R^2^>0.85, we determined the appropriate β values with the “pickSoftThreshold” function in WGCNA to emphasize strong associations and penalize weak associations between the four groups and gene expression. Based on analysis of gene significance and module membership, the co-expressed genes were clustered into modules, and connections between groups and modules were established. Functional enrichment was performed using the Metascape website and “fgsea” package. Metascape is a useful resource in gene function annotation research that provides access to multiple independent knowledge bases in one portal, including functional enrichment analysis and gene annotation tools [29]. Subsequently, we merged two groups that showed neither the best nor the worst survival into a “mixed” group, and we used the “limma” package to identify DEGs among the three resulting groups. After acquiring the DEGs, we performed fast gene set enrichment analysis using the “fgsea” package (https://github.com/ctlab/fgsea/) with the following criteria for statistically significant biological processes: p<0.1 and absolute normalized enrichment score>1 [30].

### 2.5 Tracking tumor immunophenotype (TIP) analysis

TIP analysis was used to investigate the tumor immune cycle and evaluate seven anti-cancer immune stages, and to visualize the proportion of cells in each immune stage. From each group, ten random samples were extracted for TIP analysis using the website http://biocc.hrbmu.edu.cn/TIP/ [31].

### 2.6 Combined score classifier and external validation

Survival analysis was conducted on the new (merged) groups. Then, GLC-TME score was evaluated with respect to its potential as an independent prognostic factor in breast cancer. Age, gender, tumor stage and GLC-TME score were identified as potential prognostic factors, and multivariate Cox analysis was performed to investigate the risk associated with each factor in terms of the survival of breast cancer patients. The METABRIC and GSE42568 datasets were used for validation of the key prognostic genes, and a prognostic model was established. After normalization of the datasets, the expression of key prognostic genes was analyzed in breast cancer and control groups.

### 2.7 Quantitative real-time polymerase chain reaction (qRT-PCR)

The relative expression of genes in the MCF-7 and MCF-10A cell lines was determined using qRT-PCR. Total RNA of cells was extracted with an RNA isolation kit (Accurate Biology, catalog# AG21023), and HiScript III RT SuperMix for qPCR (+gDNA wiper) Kit (Vazyme, catalog# R323– 01) was used for the reverse transcription reaction. qRT-PCR was then performed to quantitively identify gene expression with ChamQ Universal SYBR qPCR Master Mix (Vazyme, catalog# Q711– 02). The specific primer sequences used are presented in Supplementary Table 1.

### 2.8 Cell culture

Luminal breast cancer cell line MCF-7 and normal breast epithelial cell line MCF-10A were acquired from ATCC (Manassas, Virginia, USA). Cells were cultured in Dulbecco’s modified Eagle’s medium (KeyGEN BioTECH, Nanjing, China) supplemented with 10% fetal bovine serum (FBS, LONSERA), 80 U/ml penicillin, and 0.08 mg/ml streptomycin and incubated at 37 °C in 5% CO_2_.

### 2.9 Cell transfection

To perform knockdown of selected genes, specific short interfering RNAs (siRNAs) were synthesized by Biotechnologies (Nantong, China); the sequences are provided in Supplementary Table 2. Transfection was performed when the cells reached 60–80% confluence, according to the instructions provided with jetprimer (Polypuls, catalog# 101111146). The siRNA was diluted with 200 μL buffer, and the mixture was vortexed; then, 4 μL jetprimer was added, following by vortexing for a further 10 s. The transfection mixture was allowed to stand for 10–15 min at room temperature before being added to the culture medium (200 μl/well).

### 2.10 Wound healing assay

To determine the migration ability of tumor cells, a wound-healing assay was carried out. A wound was created on the cell surface; then, the cells were washed once with phosphate-buffered saline and cultured in a low serum concentration (≤2%). At two timepoints (24 h and 48 h), the wound was observed via a microscope. The migration rate was calculated using the following formula: migration (%)=(D_t_□−□D_0_) /D_0_, where Dt represents the width of the wound at time t, and D_0_ is the initial width.

### 2.11 Transwell invasion assay

For the invasion assay, 1 × 10^5^ MCF-7 cells suspended in 200 μL serum-free medium were plated on BD BioCoat Matrigel Invasion Chambers (pore size 8 μm; BD Biosciences), and the lower chamber was filled with medium containing 20% FBS. After 48 h of culture, invasive cells were stained with crystal violet solution according to the standard protocol. Cell numbers were counted in five random fields using phase contrast microscopy.

### 2.12 Cell counting kit-8 (CCK8) assay

To investigate the proliferation ability of cells, a CCK8 assay was performed. Cells were seeded in 96-well plates at a density of 3,000 cells/well and cultured at 37 °C, 5% CO_2_, for 1 day, 2 days, or 3 days. Numbers of live cells were identified using CCK8 reagent (10 μl/well; Beyotime, Shanghai, China), and the absorbance was detected at 450 nm with a microplate reader (Bio-Rad iMark).

### 2.13 Colony formation assay

Cells were seeded in six-well plates at a density of 1,000 cells/well and cultivated at 37 °C. The culture medium was replaced with fresh medium every 3 days. All colonies were stained with 0.1% crystal violet at room temperature after 5 days of culture. After staining, the colonies were observed and counted.

### 2.14 RNA-seq and transcriptomic analysis

Cells were harvested and total RNA was extracted using an RNA mini kit (Qiagen, Germany). Enrichment of mRNA, fragmentation, reverse transcription, library construction, sequencing (Illumina NovaSeq 6000), and data analysis were performed by Genergy, Biotechnology Co. Ltd. (Shanghai, China). The acquired raw reads were handled with Skewer and checked for quality using FastQC. Clean reads were aligned to human genome hg38 using STAR and then quantified with StringTie. The expression count matrix was analyzed with package DESeq2 (v. 1.38.3) for DEG identification, with the criterion p<0.05. To evaluate the glucose metabolism activity of samples, gene set variation analysis (GSVA) scores were calculated using the GSVA (v. 1.46.0) package in gene set “GOBP_GLUCOSE_METABOLIC_PROCESS”. Expression of significant hub genes at different breast cancer stages was determined using the GEPIA database (http://gepia2.cancer-pku.cn/#analysis).

The “bulkPseudotime” package (https://github.com/junjunlab/bulkPseudotime) was used for pseudo-time analysis of sequenced samples. Briefly, principal component analysis (PCA) was applied to bulk-seq data for specific genes. The pseudotime was calculated based on the distance between neighboring samples and was scaled to a range of 0 to 10. For each gene, the expression model was fit based on expression level and pseudotime using the “loess” function in R, and 500 time points were generated between 0 and 10. The corresponding expression levels were calculated based on the fitted curve and normalized by z-score.

### 2.15 Single-cell data processing and cell type identification

The “Seurat” package (v. 4.3.0) was used for single-cell dataset processing [32]. From GSE195861, 13 breast cancer samples and one normal breast sample were extracted for analysis. Low-quality cells were removed using the following screening criteria: 200<nFeature_RNA<4,000 and percentage.mt<15%. The results were normalized using the “LogNormalize” function with the default parameter (scale factor=10,000), and PCA was performed. The top ten principal components were selected for removal of batch effects with the “Harmony” package (v. 0.1.1) [33], and Uniform Manifold Approximation and Projection dimension reduction was used for cell clustering. Cluster annotation was based on canonical markers reported in previous studies.

After cell annotation, the expression distributions of significant hub genes were visualized using the “schex” (https://github.com/SaskiaFreytag/schex) and “Scillus” (https://github.com/xmc811/Scillus) packages. To identify pathologic cell types of breast cancer, we used the scPagwas algorithm to integrate breast cancer GWAS dataset ukb-b-16890 from the IEU database (https://gwas.mrcieu.ac.uk/datasets/ukb-b-16890/) with an annotated single-cell dataset [34]. A trait-relevant score for each cell was computed by averaging the expression levels of the trait-relevant genes and subtracting the random control cell score via the cell-scoring method used in Seurat.

### 2.16 Exploration of luminal cells in breast cancer

Through cell annotation and scPagwas analysis, we identified luminal cell clusters as the pathologic cell type in breast cancer. Therefore, we extracted all luminal cells from samples and performed re-clustering. For pseudo-time analysis, we used the “Monocle” package (v. 2.26.0) [35]. Re-clustered luminal cells were screened with the following criteria: mean expression>0.1 and dispersion empirical>1×dispersion fit cells. Dimension reduction was performed by the “DDRTree” method, and specific cell differentiation trajectories were identified with the “reduceDimension” function.

Particularly, we downloaded the cancer cell stemness gene set “MALTA_CURATED_STEMNESS_MARKERS.v2024.1.Hs.gmt” from MsigDB database, and calculated the used the “AddModuleScore” function to calculate the cell stemness scores in luminal subclusters. The subcluster with highest cell stemness score was assigned as the root node of the trajectory.

Moreover, we calculate the gene score of significant hub genes (PGK1, SIRT7, PMAIP1 and SORB1) using “AddModuleScore” function. After scores had been calculated for each cell, the luminal cells were divided into Score-high and Score-low groups. The characteristic genes in the different luminal clusters were identified using the “FindAllMarkers” function, and functional enrichment analysis was conducted.

### 2.17 Cell–cell communication and metabolic activity evaluation

Focusing on the gene activation or not in luminal cells, we divided luminal cells into PMAIP1+ and PMAIP1− groups according to the expression or non-expression of the gene. Then, 2,000 random tumor cells were extracted from tumor samples for cell–cell communication analysis and metabolism pathway analysis. The “Cellchat” package (v. 1.6.1) was used to analyze cell-state specific signaling communications between cell clusters in samples [36]. Intercellular interactions between ligand and receptor activities were identified based on prior knowledge, and probabilities of interactions between cells were then inferred by random permutations. The inferred communication probability was visualized using a weighted directed graph representing the intercellular communication network.

The metabolic pathway activities in cells of different types in breast cancer were investigated using the “scMetabolism” package (v. 0.2.1) [37], which includes 85 Kyoto Encyclopedia of Genes and Genomes (KEGG) and 82 Reactome metabolic-related pathway entries. The different types of cells in single-cell samples were evaluated and scored based on the conventional single-cell matrix file.

### 2.18 Bayes deconvolution of single-cell data

To evaluate the cell composition of TCGA-BRCA bulk-seq samples, we used the BayesPrism (https://github.com/Danko-Lab/BayesPrism) package [38], which models a prior from cell-type-specific expression profiles from single-cell RNA-seq data to jointly estimate the posterior distribution of cell type composition and cell-type-specific gene expression from bulk RNA-seq expression data of tumor or non-tumor samples. The outlier genes in the single-cell RNA-seq and bulk-seq data were removed, and the protein-encoding genes in the two datasets were selected as the BayesPrism input, with the outlier.cut threshold set to 0.01 and the outlier.fraction threshold set to 0.1. Subsequently, the BayesPrism process was executed, and the calculated final Gibbs theta values were used as estimates for the fraction of each type of cells in each sample of the TCGA-BRCA dataset.

After acquiring the cell composition of each sample, the overall survival rates of patients with different PMAIP1+ luminal and PMAIP1− luminal cell proportions were analyzed using the “survival” package. Specifically, proportion ratios were calculated using the formula *proportion ratio - prop(PMA/P1+ luminal cells)/prop(PMA/P1-luminal cells)*. An optimized cutoff of proportion ratio was calculated with the “surv_cutpoint” function. The proportions of PMAIP1+ luminal cells in patients with different clinical subtypes were also explored.

According to the median PMAIP1+ luminal cell proportion, patients were divided into high and low PMAIP1+ luminal cell groups. Immune-related gene sets were downloaded from the MSigDB, and the expression of immune-related genes in the high and low PMAIP1+ luminal groups was explored.

### 2.19 Tumor phenotype analysis of deconvoluted bulk-seq samples

We downloaded single nucleotide variation data of breast cancer patients from TCGA. Tumor mutation load and gene mutations of each sample were obtained using the maftools (v. 2.14.0) package in R, and immune escape and dysfunction in tumors were evaluated using the “Tumor Immune Dysfunction and Exclusion (TIDE)” database (http://tide.dfci.harvard.edu/). TIDE evaluates the potential clinical efficacy of immunotherapy in different risk groups, reflecting the potential ability for tumor immune escape; higher TIDE scores are associated with poor immunotherapy efficacy.

We also predicted the drug sensitivity of the high and low PMAIP1+ luminal cell groups using the “oncoPredict” package [39], by calculating half-maximal inhibitory concentration values of commonly used drugs in TCGA-BRCA samples based on the Genomics of Drug Sensitivity in Cancer database (https://www.cancerrxgene.org/).

Furthermore, the WGCNA method was used to identify co-expression modules that were related to PMAIP1+ luminal cells in the TCGA-BRCA dataset. The gene module that was most strongly correlated with PMAIP1+ luminal cells was extracted and intersected with DEGs identified in the cell-line sequencing data. A protein–protein interaction network was constructed using the STRING database (https://string-db.org/), and KEGG and gene ontology functional enrichment analyses were subsequently conducted.

### 2.20 Statistical analysis

Statistical analyses were performed using R v. 4.3.0 (http://www.rproject.org). In the bioinformatic analyses, log-rank test was used for calculations of significance in the survival analysis, and Wilcoxon test was used to compare differences. Original experimental data were processed and analyzed with GraphPad Prism 9 and are presented as mean ± s.e.m. Multiple t-tests were used to calculate p-values; *p<0.05, **p<0.01, ***p<0.001, ****p<0.0001.

## 3 Results

### 3.1 Significant glucose-metabolism-related genes in breast cancer

The R packages “limma” and “sva” were applied to identify DEGs from glucose metabolism-related geneset in breast cancer. A total of 151 DEGs were identified with a threshold of adjusted p<0.05 (Figure 1A). Then, univariate Cox regression analysis of the DEGs in breast cancer was performed. Fourteen prognosis-related DEGs were identified, of which four key prognostic genes were screened using the LASSO algorithm from “glmnet” package (Figure 1B). These four genes were subjected to multivariate Cox analysis to evaluate their prognostic value (Figure 1C) and subsequent GLC-Score calculation. Survival analysis of breast cancer patients based on GLC-Score showed significant differences between the survival probabilities of the high-score group and low-score group, with higher scores associated with survival probability (Figure 1D).

**FIGURE 1.**
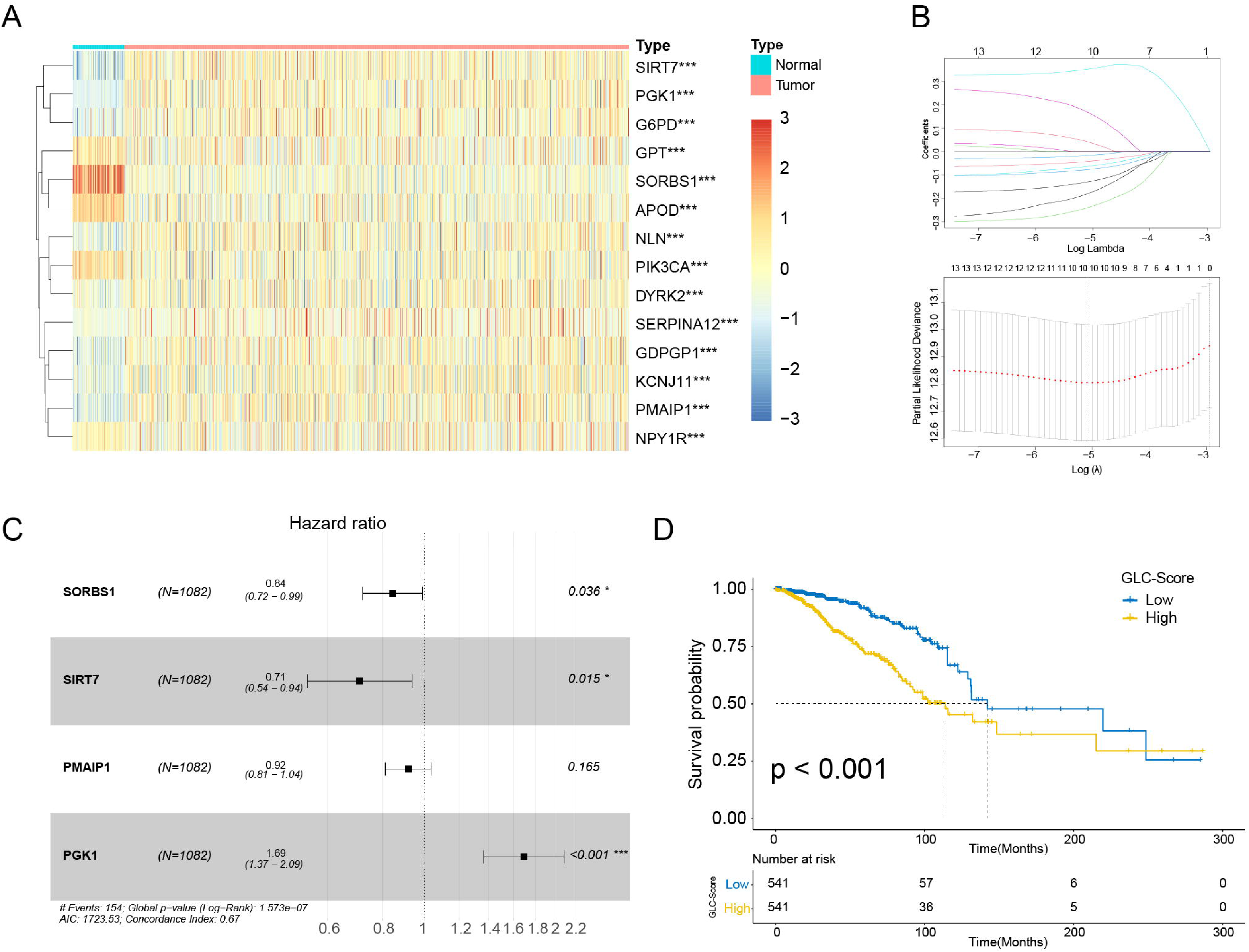
The identification of key glucose metabolism genes in breast cancer and relevant prognosis analysis. (A) Heatmap revealing expression differentiation of glucose metabolism genes selected by univariate COX regression in the TCGA-BRCA cohorts, with the threshold of adj.p<0.05. (B) Key prognostic genes screened using LASSO algorithm. (C) The prognosis analysis of key prognostic genes conducted by multivariate COX. (D) Kaplan-Meier survival curves of breast cancer patients in high and low GLC-Score group.

### 3.2 Immune infiltration analysis and protective immune cell identification

Immune-related cells have significant roles in the formation of the TME; thus, immune infiltration analysis can be used to evaluate the TME. Here, we used CIBERSORT for estimation of immune infiltration in breast cancer samples; the relative immune cell proportions calculated in this way are presented in Figure 2A. The effects of different types of immune cells on survival of breast cancer patients were analyzed with a threshold of p<0.05 (Figure S1). An optimal cutoff of the cell proportion value was determined using the “surv_cutpoint” function, and patients were divided into high and low cell proportion groups. The immune cells that had significant effects on survival of breast cancer patients included CD8 T cells, activated T cells, and M1 macrophages (Figure 2B).

**FIGURE 2.**
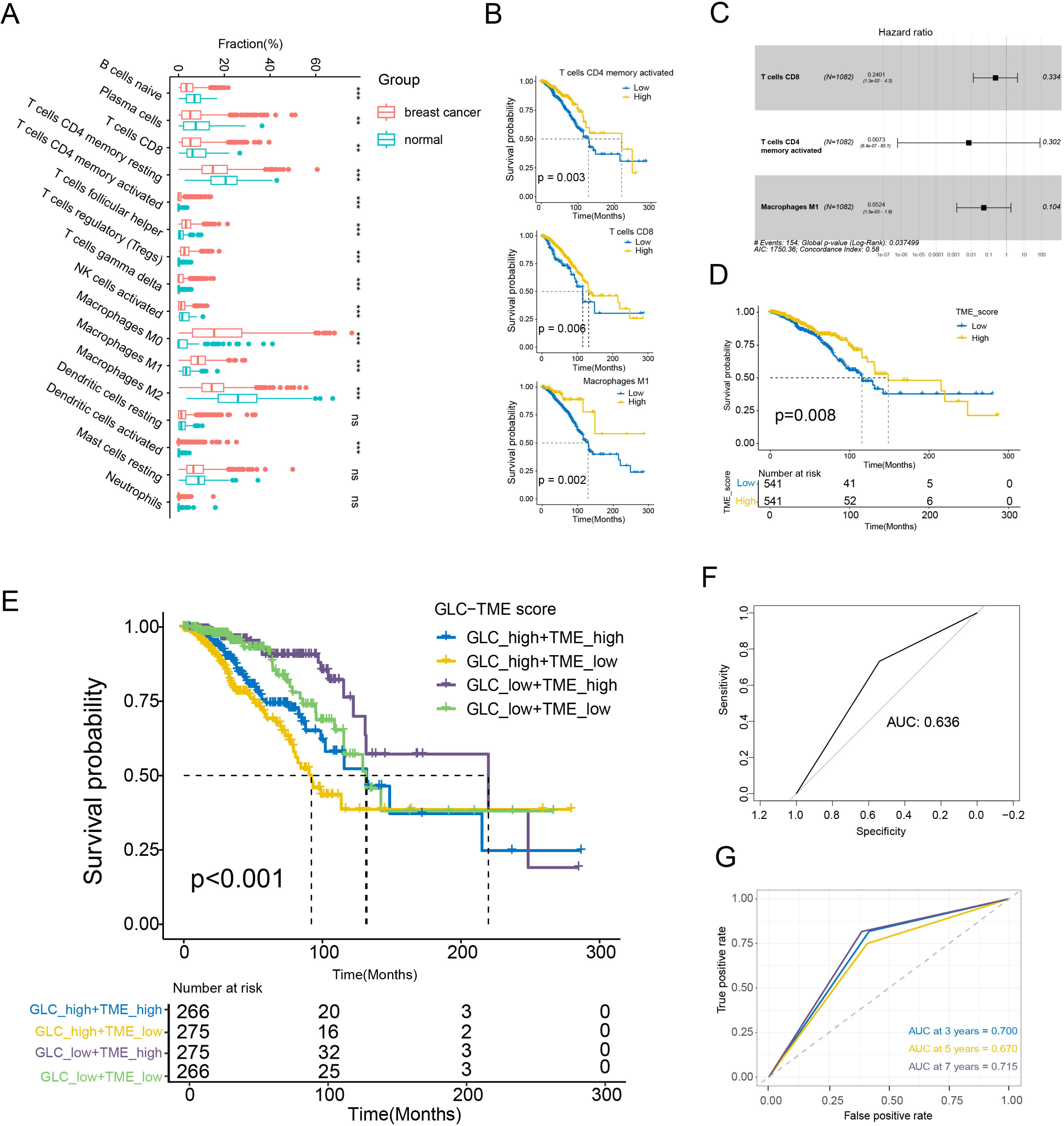
Immune infiltration analysis and prognostic model establishment. (A) Comparison of different immune cells’ ratios in breast cancer and control groups. (B) The survival analysis of selected protective immune cells. (C) Multivariate COX regression of selected immune cells. (D) Kaplan-Meier survival curves of breast cancer patients in high and low TME-Score group. (E) The establishment of prognostic model combining GLC-Score and TME-Score. (F) The ROC curve of the GLC-TME prognostic model. (G) The ROC curves of predictive ability for predicting 3-, 5-, 7-year survival of breast cancer patients via GLC-TME prognostic model.

Multivariate Cox analysis was then conducted to evaluate the prognostic value of these immune cell types in patients with breast cancer (Figure 2C). TME-Score was calculated via the formula 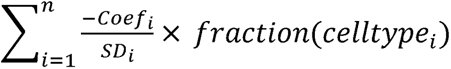. Survival analysis of breast cancer patients based on TME-Score showed significant differences in survival between the high-score group and low-score group, with lower scores being correlated with lower survival probability (Figure 2D). Patients with lower levels of immune infiltration also tended to have a poorer prognosis, supporting the predictive ability of TME-Score.

### 3.3 Establishment of prognostic model

We next combined the GLC-Score and TME-Score to establish a GLC-TME prognostic model with two dimensions for evaluation. Breast cancer patients were classified into four groups on the basis of the two scores (GLC-High+TME-High, GLC-High+TME-Low, GLC-Low+TME-High, and GLC-Low+TME-Low), and survival analysis was performed on these groups (Figure 2E). The GLC-Low+TME-High group had the best survival, whereas the GLC-High+TME-Low group had the poorest. Receiver operating characteristic (ROC) curves were used to measure the model’s specificity, for which the area under the curve was 0.636 (Figure 2F). Moreover, the area under the curve values for 3-year, 5-year, and 7-year prediction were 0.700, 0.670, and 0.715 respectively (Figure 2G), demonstrating that the predictive ability of the prognostic model improves with increasing overall survival time.

### 3.4 Identification of co-expressed genes and functional enrichment

We used WGCNA to identify co-expressed genes in the TCGA-BRCA cohort, so that the major functional pathways in breast cancer could be explored. To construct a scale-free co-expression network, a soft threshold of 6 (R^2^=0.87) was picked. The clustering dendrogram of samples is illustrated in Figure 3A. The 13 gene co-expression modules were generated in different colors, and the correlations among the four patient groups and the co-expression modules are presented in Figure 3B. The green module showed the strongest correlation with the GLC-High+TME-Low group, whereas the red module showed the strongest correlation with the GLC-Low+TME-High group.

**FIGURE 3.**
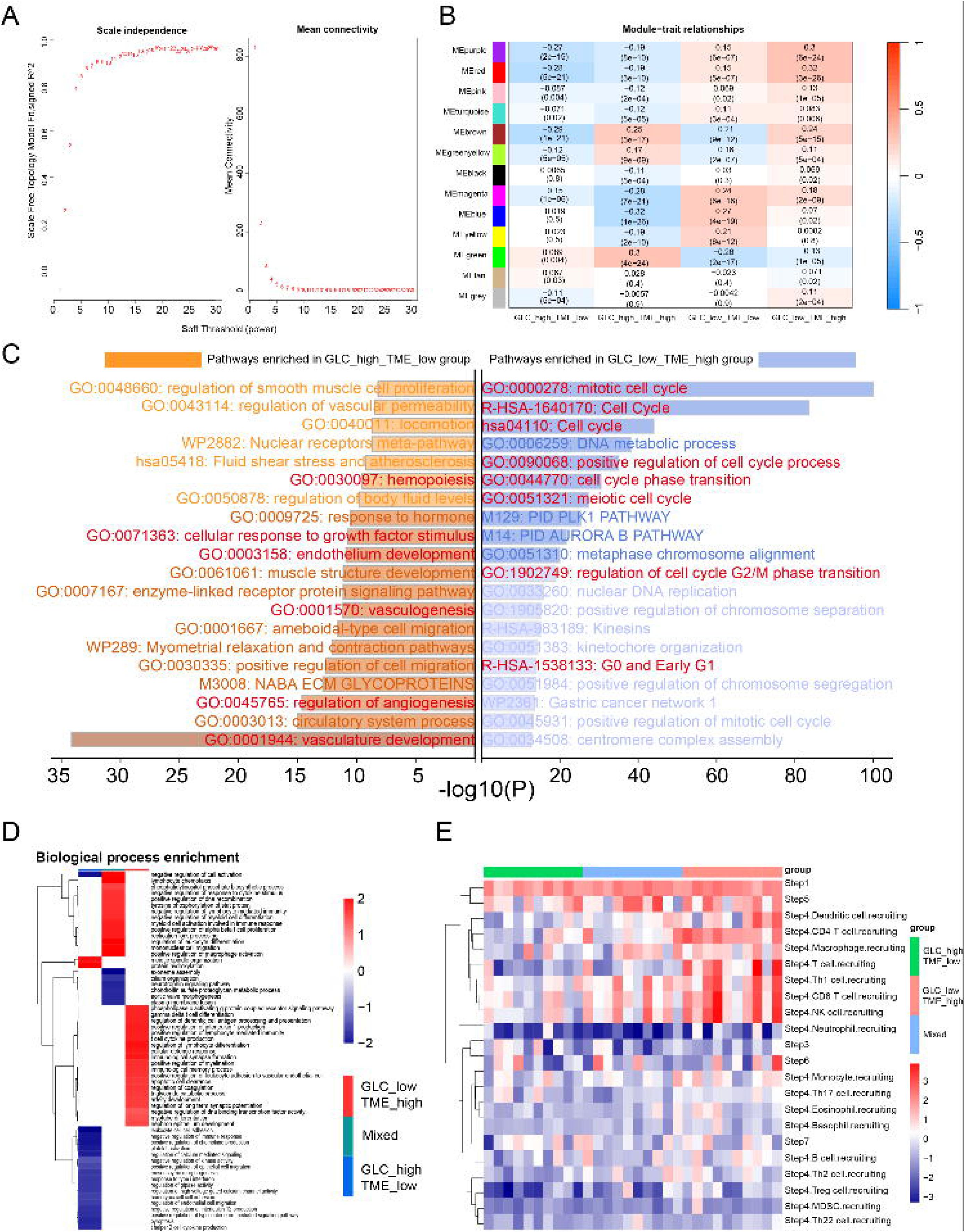
Co-expressed gene identification, functional enrichment and TIP analysis of different groups in breast cancer. (A) β=6 is chosen to be the soft threshold with the co-analysis of scale independence and average connectivity. (B) Heatmap of the association between modules and groups. The correlation coefficients and p-values were presented in blocks respectively. (C) Enriched signaling pathways of GLC-High+TME-Low and GLC-Low+TME-High groups via metascape. Significant pathways were marked red. (D) Differentially enriched signaling pathways between GLC-Low+TME-High, Mixed and GLC-High+TME-Low groups using fgsea. (E) TIP analysis of three groups.

Functional enrichment analysis of the co-expressed genes via the Metascape portal indicated that pathways related to vascular development and circular system were the most enriched in the GLC-High+TME-Low group, whereas pathways related to the cell cycle and DNA metabolic process were the most enriched in the GLC-Low+TME-High group (Figure 3C). The enriched pathways showed upregulated expression of genes related to vascular development in GLC-High+TME-Low group, providing an explanation for the poor overall survival of patients in this group.

As the GLC-High+TME-High and GLC-Low+TME-Low groups showed neither the best nor the worst survival, they were merged into a “Mixed” group. Fast gene set enrichment analysis revealed differences in pathway activities among the three resulting groups. The GLC-Low+TME-High group showed significant increased activity in of T cell-related and immune-related pathways, whereas the GLC-High+TME-Low group showed decreased activity of pyroptosis and immune-related pathways (Figure 3D). TIP analysis of the three groups indicated that the most immune cells were recruited in the GLC-Low+TME-High group, whereas in the GLC-High+TME-Low group, the recruitment of immune-related cells was deactivated. The Mixed group showed a transition state between activated recruitment and non-recruitment of immune cells (Figure 3E).

### 3.5 External validation of significant glucose metabolism genes

After merging the GLC-High+TME-High and GLC-Low+TME-Low groups, we performed survival analysis. The GLC-Low+TME-High group still showed the best survival, whereas the GLC-High+TME-Low group showed the worst (Figure 4A). To evaluate the individual prognostic value of the GLC-TME Score, multivariate Cox analysis was conducted. The results indicated that the GLC-TME Classifier is capable of functioning as a predictor of breast cancer risk, as its hazard ratio was 1.68 (95% confidence interval: 1.315–2.2), with a significant p<0.001 (Figure 4B). The METABRIC dataset was used to validate the performance of the prognostic model in clinical practice; this showed that the GLC-TME prognostic model was robust and reliable for a general background (Figure S2). In addition, survival analyses were performed involving patients with specific clinical subtypes; the GLC-Low+TME-High group maintained the best survival among patients of major clinical subtypes, including those aged <65 years and those with different tumor stages, whereas the GLC-High+TME-Low showed the worst survival (Figure S3A). Based on the identified independent risk factors for breast cancer, a nomogram for survival prediction was constructed (Figure S3B). Systematic analysis of clinical data revealed that the GLC-TME Classifier was clinically meaningful, as it was able to accurately categorize patients and predict their survival.

**FIGURE 4.**
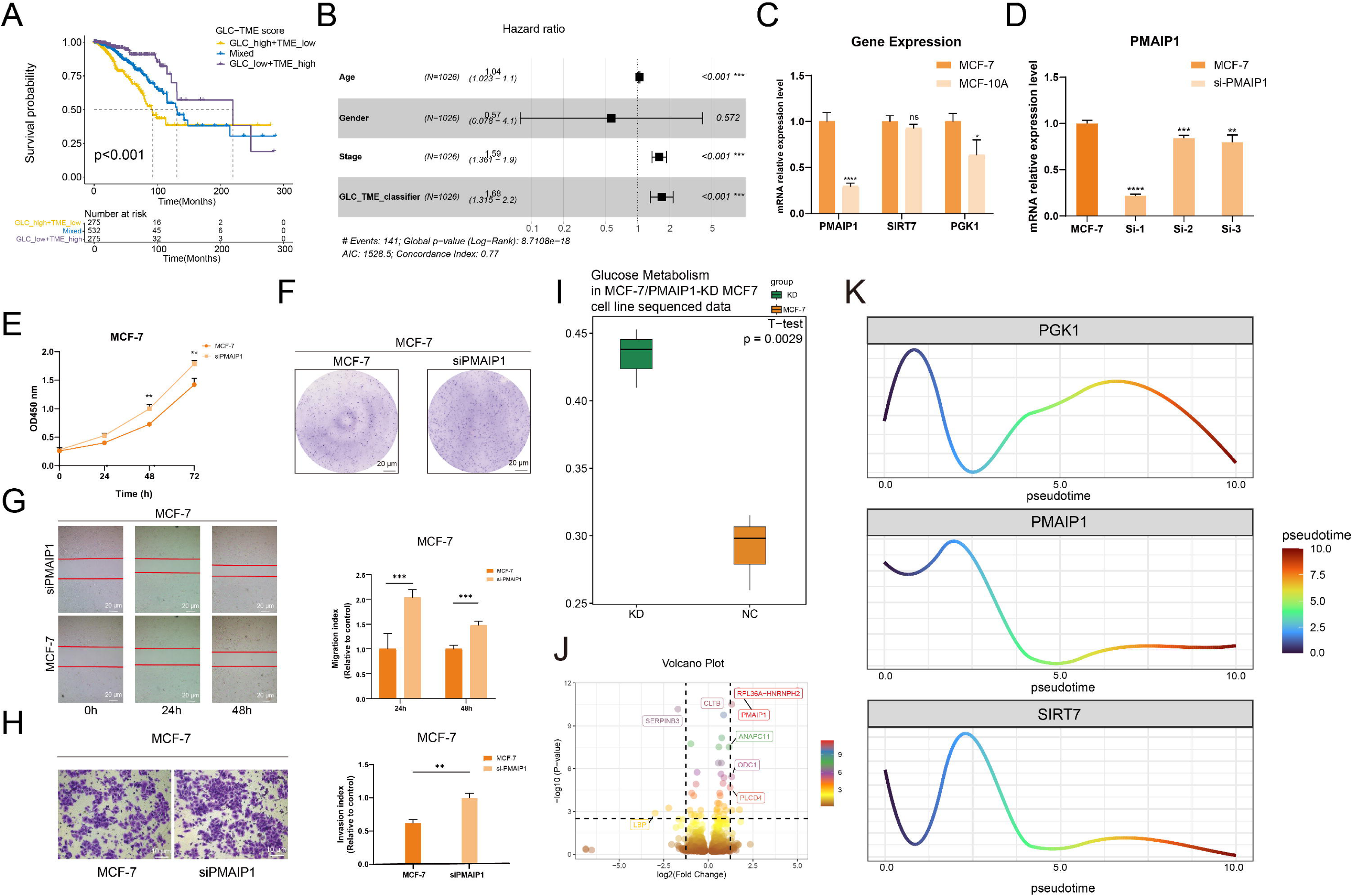
External validation and the exploration of PMAIP1 in breast cancer. (A) The Kaplan-Meier survival curve of GLC-TME Score after merging groups. (B) Multivariate COX regression of clinical risk factors in breast cancer, including age, gender, tumor stage and GLC-TME Score classifier. (C) qPCR analysis of PMAIP1, PGK1, and SIRT7 expression in MCF-7 and MCF-10A cells. (D) qPCR analysis of PMAIP1 knockdown efficiency. (E) CCK-8 experiment validates the increased proliferative capacity of MCF-7 cells after PMAIP1 knockdown. (F) Knockdown of PMAIP1 enhances colony formation in MCF-7 cells. Scale bars: 20 μm. (G) Wound healing assay reveals that PMAIP1 knockdown facilitates MCF-7 cell migration. Scale bars: 20 μm. (H) Transwell assay reveals enhanced invasion of MCF-7 cells upon PMAIP1 knockdown. Scale bars: 10 μm. (I) The GSVA scores of geneset “GOBP_GLUCOSE_METABOLIC_PROCESS” in sequenced mRNA data. (J) Volcano plot of DEGs based on bulk RNA sequencing, genes with |logFC|>1 were labelled in the plot. (K) The pseudo-time analysis conducted in MCF-7/PMAIP1-KD MCF-7 cell line sequenced samples. PMAIP1-KD MCF-7 cell line samples and MCF-7 cell line samples were used to simulate the development progression of breast cancer. Curve plots showed the expression trajectory of each hub gene and the differentiation trajectory of samples. All data are presented in mean ± sd. p values were calculated using multiple t-tests. *p<0.05, **p<0.01, ***p<0.001, ****p<0.0001.

Focusing on the significant genes, we subsequently validated the expression levels of PMAIP1, SIRT7, PGK1, and SORBS1 in external datasets, including METABRIC and GSE42568 (Figure S3D). PMAIP1, SIRT7, and PGK1 showed differential expression levels between the two datasets. Subsequently, we performed qRT-PCR experiments to further examine the expression levels of these genes in breast cancer (MCF-7) cells and breast normal epithelial (MCF-10A) cells (Figure 4C). The results showed that PMAIP1 was the most significantly upregulated glucose metabolism-related gene in MCF-7 cells. PMAIP1 is a member of the BCL-2 family, which has pro-apoptotic functions intersecting with the p53 pathway [40]. The upregulation of PMAIP1 might indicate a particular role in breast cancer.

### 3.6 PMAIP1 suppresses the proliferation, migration, and invasion of MCF-7 cells

After identifying PMAIP1 as a key prognostic gene, we constructed an siRNA (Figure 4D) to functionally knock down PMAIP1 in MCF-7 cells. Colony formation and CCK-8 experiments revealed that knockdown of PMAIP1 significantly enhanced the proliferation ability of the MCF-7 cells (Figure 4E-F). A wound healing assay demonstrated that downregulation of PMAIP1 significantly promoted migration of the tumor cells (Figure 4G), and a transwell assay showed that the invasion of MCF-7 cells was promoted by PMAIP1 knockdown (Figure 4H). These results suggest that PMAIP1 inhibits the proliferation, migration, and invasion of MCF-7 cells; thus, it is a potential oncogene in breast cancer.

Sequencing data for the MCF-7 and PMAIP-knockdown MCF-7 cell lines showed that the GSVA score of “GOBP_GLUCOSE_METABOLIC_PROCESS” in the PMAIP1-knockdown MCF-7 group was significantly higher than that of the MCF-7 group, indicating that glucose metabolism was activated in breast cancer after PMAIP1 knockdown (Figure 4I and S3C). The DEGs between the two groups were also identified using the “DESeq2” package (Figure 4J). According to the selection criteria, 1,213 DEGs were screened, comprising 636 upregulated and 577 downregulated genes. After PCA calculation and trajectory exploration, bulk-pseudotime analysis revealed variations in expression of PGK1, PMAIP1, and SIRT7 in the development of breast cancer (Figure S3E-F and 4K). The expression of the three genes at different tumor stages was also investigated via the breast cancer cohorts in the GEPIA database (Figure S3G). Integrating the results of the bulk-pseudotime analysis and the expression variation data acquired from GEPIA, we observed a decreasing trend in expression of PMAIP1, with relative low expression in the middle stage compared with the terminal stage. This expression pattern of PMAIP1 could be attributed to variation among different types of cells in breast cancer as the disease develops. Therefore, we next explored the cell heterogeneity of breast cancer at single-cell resolution.

### 3.7 Single-cell data processing

To further investigate significant glucose metabolism genes in breast cancer, we conducted single-cell analysis. After quality control, 24,241 cells were obtained for downstream analysis, and normalization and PCA were performed (Figure S4A-C). Ten principal components were selected for “Harmony” integration, which was applied for removal of batch effects (Figure S4D). Twenty-five cell clusters were identified and annotated based on canonical markers (Table 1 and Figure S4E); luminal cells, T cells, macrophages, basal cells, B cells, plasma cells, erythrocytes, fibroblasts, proliferating luminal cells, proliferating T cells were identified (Figure 5A); and the expression distributions of four significant genes in tumor samples were visualized (Figure 5B and Figure S4F). Further, we identified that luminal cells showed relatively high trait-relevant scores compared with the other main cell types using scgwas method based on dataset ukb-b-16890 (Figure 5C). Thus, luminal cells were identified as the pathologic cells in breast cancer.

**FIGURE 5.**
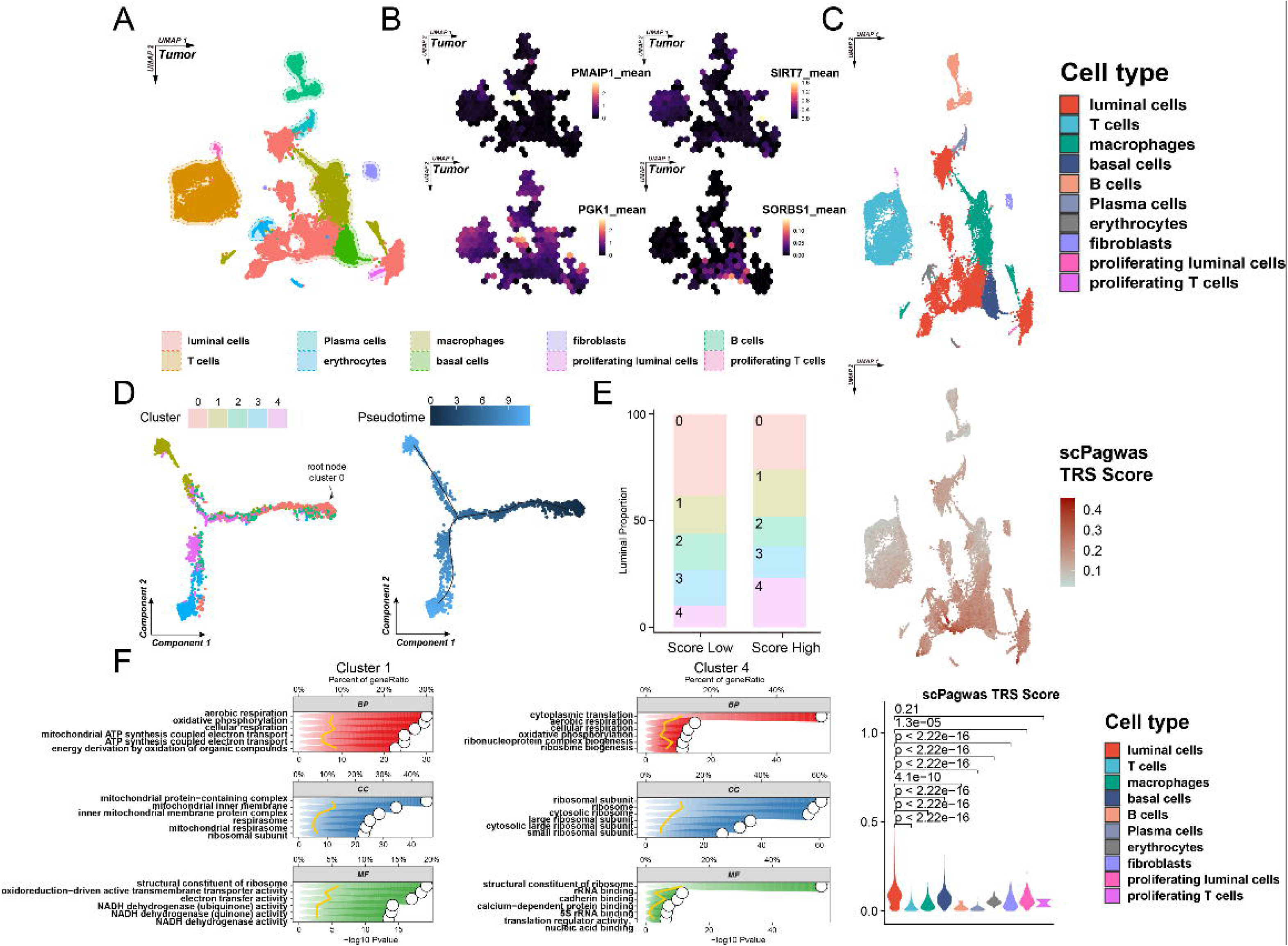
Single-cell analysis of single-cell dataset GSE195861. (A) Annotated tumor cell clusters from GSE195861. (B) Expression distribution of PMAIP1, PGK1, SIRT7 and SORBS1 in different cell types. (C) ScPagwas TRS score of different cell clusters. Luminal cells showed the highest trait relevance with the breast cancer. (D) The pseudotime analysis conducted by “monocle”. The luminal cells were re-clustered, and dark blue points to light blue points represent cell differentiation process from the earlier stage to the later stage, indicating the differentiation order. (E) The proportion of each luminal clusters in Score-Low and Score-High groups. Cluster 1 and cluster 4 showed increased proportions in high group. (F) Functional enrichment of characteristic genes in cluster 1 (left) and cluster 4 (right).

**Table. 1.**
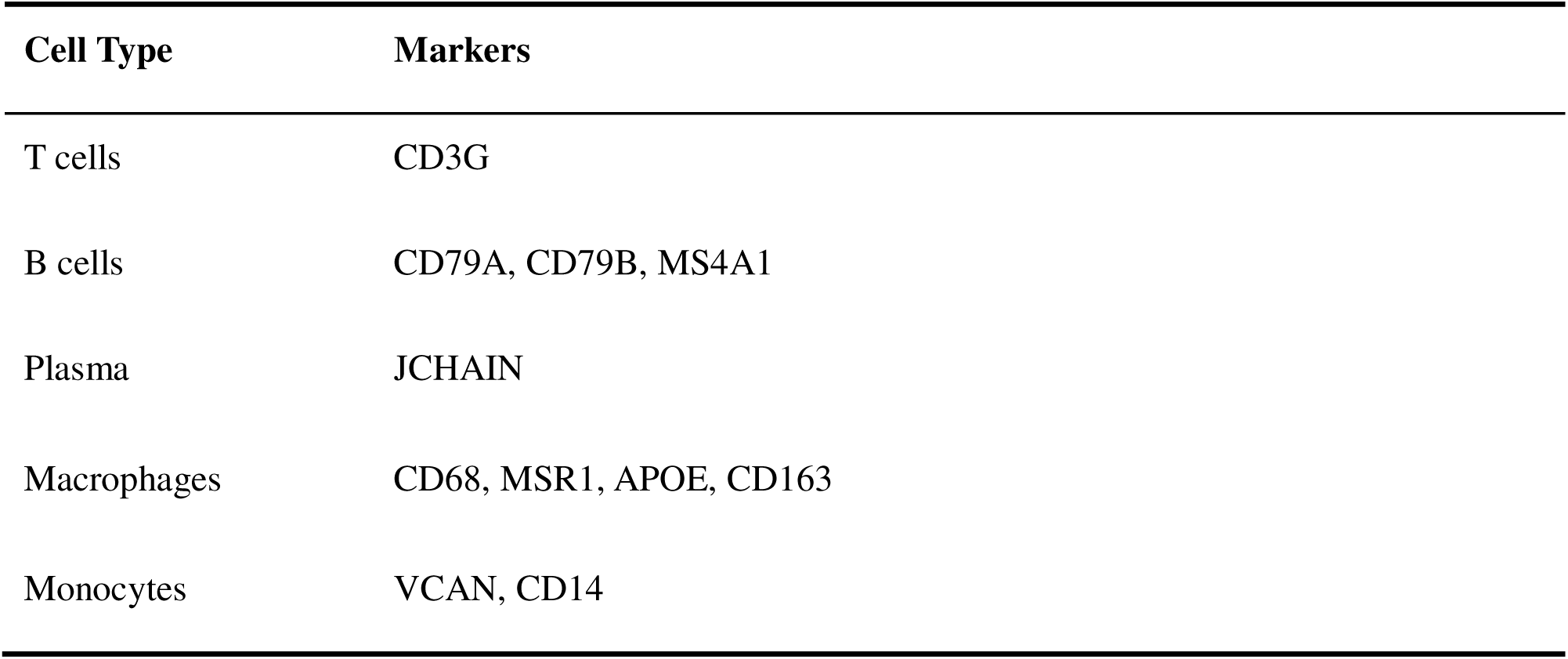
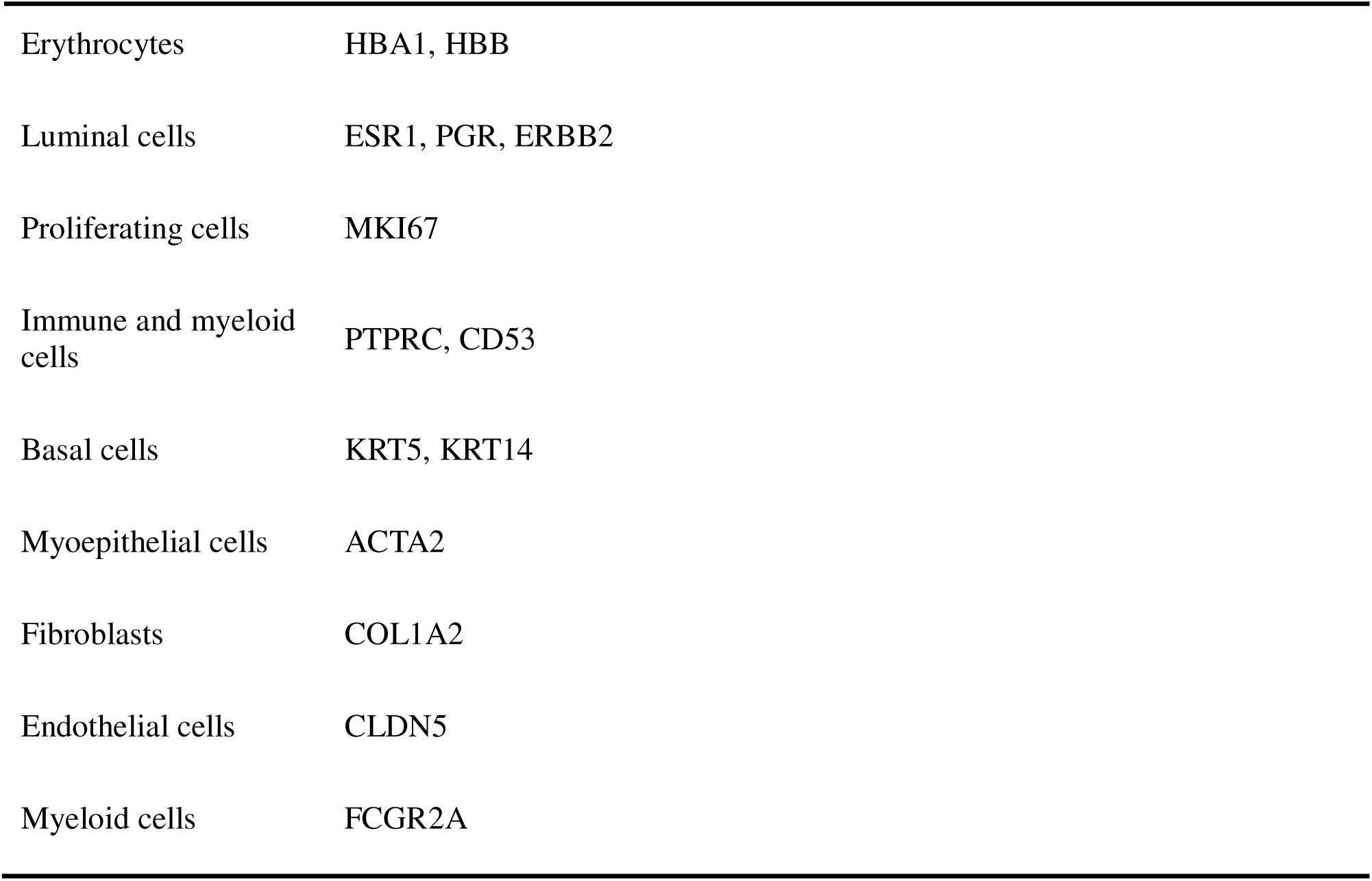
Marker genes for cell type distinction and annotation.

| Cell Type | Markers |
| --- | --- |
| T cells | CD3G |
| B cells | CD79A, CD79B, MS4A1 |
| Plasma | JCHAIN |
| Macrophages | CD68, MSR1, APOE, CD163 |
| Monocytes | VCAN, CD14 |
| Erythrocytes | HBA1, HBB |
| Luminal cells | ESR1, PGR, ERBB2 |
| Proliferating cells | MKI67 |
| Immune and myeloid cells | PTPRC, CD53 |
| Basal cells | KRT5, KRT14 |
| Myoepithelial cells | ACTA2 |
| Fibroblasts | COL1A2 |
| Endothelial cells | CLDN5 |
| Myeloid cells | FCGR2A |

After identifying the luminal clusters as the target cell clusters, luminal cells were extracted and re-clustered. The cancer cell stemness-related gene set “MALTA_CURATED_STEMNESS_MARKERS.v2024.1.Hs.gmt” was downloaded from MsigDB database, and the “AddModuleScore” function was used to evaluate cell stemness of luminal clusters (Figure S4G). Cluster 0 showed the highest stemness significantly. Higher cell stemness generally indicates earlier stage in cell differentiation [41], thus the cluster 0 was identified as the root node for pseudotime. Pseudotime analysis revealed the differentiation trajectory of luminal cell clusters (Figure 5D); The luminal clusters were observed in different areas; cluster 0 was in the initial stage of differentiation, whereas clusters 1 and 3 were in the terminal stage. To further investigate the significant hub genes in luminal differentiation, we used the “AddModuleScore” function. According to the median value of the scores calculated, luminal cells were divided into two groups: high score luminal cells and low score luminal cells. As shown in Figure 5E, clusters 1 and 4 had higher proportions of cells in the high score group compared with other clusters. Furthermore, we identified characteristic genes in clusters 1 and 4 using the “FindMarkers” function and conducted functional enrichment analysis (Figure 5F). Pathways related to cellular respiratory functions, including aerobic respiratory, oxidative phosphorylation, and cellular respiratory, were the most enriched in both clusters.

### 3.8 Cell–cell communication and metabolic landscape in breast cancer

Based on the expression or non-expression of PMAIP1, the luminal cells were divided into PMAIP1+ luminal and PMAIP1− luminal cell groups. The expression difference of PMAIP1 between the PMAIP1+/- luminal cells indicated that PMAIP1 showed distinguishing difference in PMAIP1+/- luminal cells. Thus, we can infer that the activation of PMAIP1 might introduce significant heterogeneity to luminal cells (Figure S4H). We then extracted 2,000 random cells from tumor samples for cell–cell communication analysis. Using the “Cellchat” package, the interactions between different cell clusters were inferred (Figure 6A). Compared with PMAIP1− luminal cells, PMAIP1+ luminal cells showed stronger overall interactions, especially involving the migration inhibitory factor (MIF) and galectin pathways, which indicate interactions with macrophages (Figure 6B).

**FIGURE 6.**
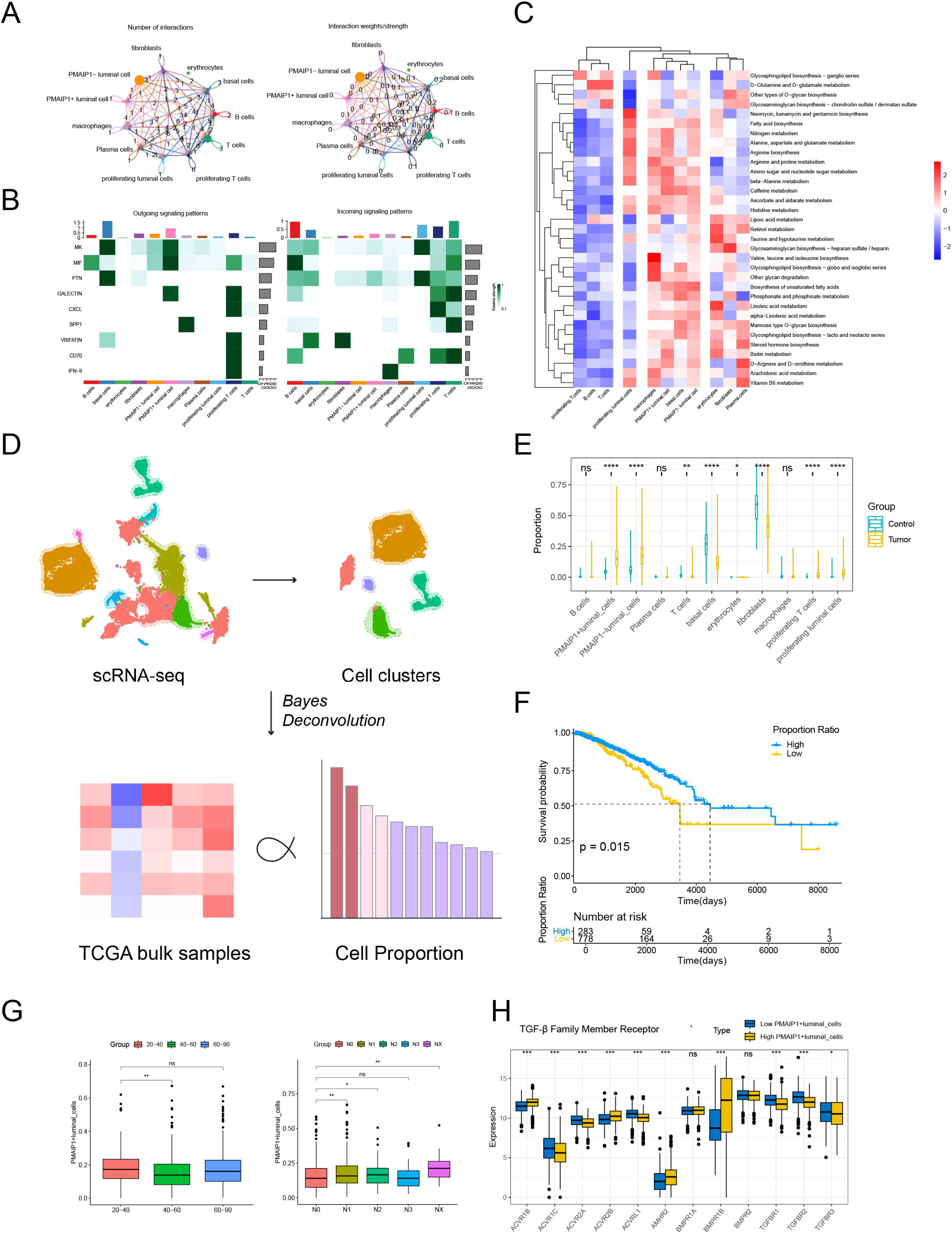
Cell-cell communication, metabolic pathway analysis and Bayes deconvolution of single-cell data. (A) Landscape of intercellular communication. The colors indicate the sender of different cell-cell communications, the edge thickness represents communication weight, and the arrow marked the communication direction. (B) Heatmap visualizing important ligands and receptors of subclusters functioning as signalers and receivers in intercellular communication activities. (C) Heatmap showing top 5 activated metabolic pathways in different tumor cell types, colors indicate the effect size. (D) Illustration of Bayes deconvolution process, mapping to TCGA-BRCA samples. (E) The proportions of different cell clusters in TCGA-BRCA samples after Bayes deconvolution process. (F) The differential survival between breast cancer patients, grouped by identified PMAIP1+ luminal proportion ratio. Patients with higher proportion ratio showed better survival outcome. (G) The proportion of PMAIP1+ luminal cells in breast cancer patients with different clinical subtypes. (H) The expression of TGF-beta family member receptors between high PMAIP1+ luminal group and low PMAIP1+ luminal group.

To explore the metabolic differences between cell clusters in breast cancer, we used the “scMetabolism” package for assessment of metabolic pathways based on the extracted 2,000 tumor cells. The top five differentiated metabolic pathways in each cell type were visualized (Figure 6C). The metabolic activity of immune cells, including T cells and B cells, declined, whereas PMAIP1− luminal cells showed increased activity in glycosylation-related pathways.

To associate the proportions of different cell clusters with bulk-seq samples, deconvolution of single-cell data was conducted. The workflow of “BayesPrism” R package was illustrated in Figure 6D. Figure 6E shows the different proportions of each cell type in tumor and control samples.

Specifically, the proportions of PMAIP1+ luminal and PMAIP1− luminal cells in tumor samples were significantly higher than those in control samples. The following proportion ratio was used for survival analysis: *proportion ratio - prop(PMA/P1+ luminal cellsJ/prop(PMA/P1-luminal cells)*. An optimal cutoff of value of the proportion ratio was determined by “surv_cutpoint” function for patient grouping, which showed differential survival outcomes (Figure 6F).

In addition, the proportion of PMAIP1+ luminal cell patients varied among patients with different clinical subtypes (Figure 6G). Patients aged 20–40 years had the highest proportion of PMAIP1+ luminal cells, with the proportion decreasing in patients aged 40–60 years. According to the median value of the PMAIP1+ luminal cell proportion, TCGA-BRCA samples were divided into a high PMAIP1+ luminal cell group and low PMAIP1+ luminal cell group. The transforming growth factor β (TGF-β) family regulates cell proliferation, growth, differentiation, and movement and is crucial to tumor proliferation and development [42]. As shown in Figure 6H, most of the TGF-β receptor genes were significantly downregulated in the high PMAIP1+ luminal cell group. TGF-β receptors activate downstream signaling pathways, inhibiting the proliferation of malignant epithelial cells. This inhibitory effect is particularly evident in normal cells and early cancer cells, helping to prevent tumor formation and growth.

### 3.9 Variation of tumor phenotypes related to PMAIP1+luminal cells

The mutational profiles of patients in the high and low PMAIP1+ luminal cell groups based on TCGA data were analyzed (Figures 7A and S5A). Generally, patients in the low PMAIP1+ luminal cell group showed increased rates of gene mutations; in particular, TP53 was mutated in 49% of patients. In the high PMAIP1+ luminal cell group, TP53 and GATA3 mutations were each found in 20% of patients. TP53 is critical to the regulation of the cell cycle, proliferation, and apoptosis. Thus, the increased rates of TP53 mutation in the low PMAIP1+ luminal cell group represent a loss of tumor suppressor functions, as well as increased tumor invasiveness and malignancy.

**FIGURE 7.**
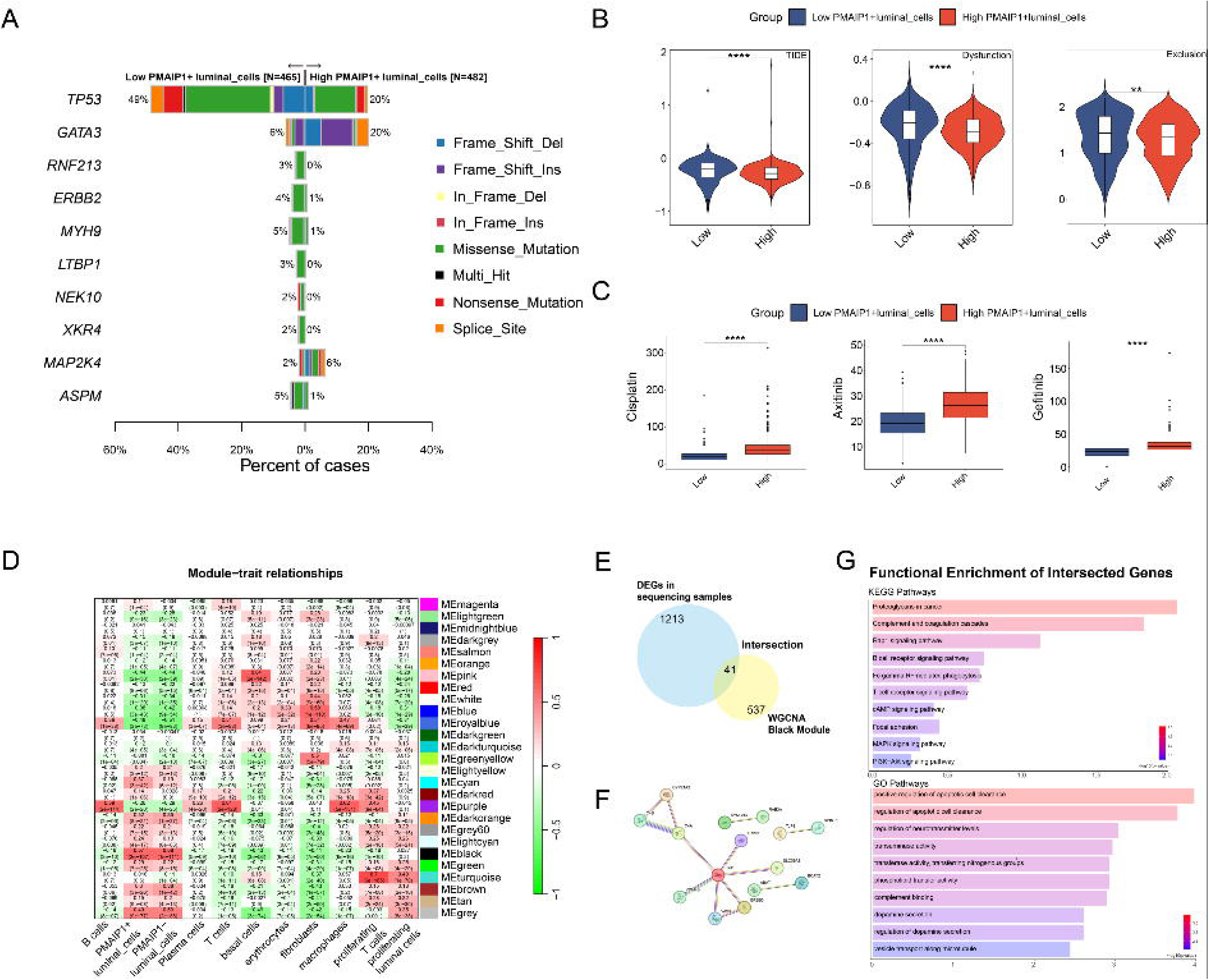
The relationships between the variations of tumor phenotypes and the proportion of PMAIP1+ luminal cells. (A) Mutation landscape of patients in high and low PMAIP1+ luminal cell groups. (B) TIDE score, dysfunction score and exclusion score in high and low PMAIP1+ luminal cell groups. (C) Drug sensitivity prediction of cisplatin, axitinib and gefitinib. (D) Heatmap of the association between modules and cell types. The correlation coefficients and p-values were presented in blocks respectively. (E) The intersection between DEGs in sequencing samples and genes in black module. (F) Protein-protein interaction network predicted by String database. (G) KEGG and GO functional enrichment of 41 intersected genes.

In addition, we used the TIDE webtool to evaluate the potential clinical effectiveness of immunotherapy according to molecular subtypes (Figure 7B). Compared with the low PMAIP1+ luminal group, the high PMAIP1+ group had lower TIDE scores, dysfunction scores, and exclusion scores, indicating better T cell infiltration, T cell function, and immune therapy response.

Interestingly, in GLC-TME groups, the GLC_Low+TME_High group showed highest proportion of high PMAIP1+ luminal patients, which was corresponding to the recruitment track of immune cells in breast cancer (Figure S5B). Moreover, according to the drug sensitivity prediction, patients in the high PMAIP1+ luminal cell group were more sensitive to different types of anti-cancer drugs, including cisplatin, axitinib, and gefitinib (Figure 7C and S6).

We used the WGCNA method to identify a module of 537 genes co-expressed in PMAIP1+ luminal cells (Figure 7D). Intersection of the DEGs acquired from sequencing samples with this co-expression module yielded 41 genes (Figure 7E). In the protein–protein interaction network constructed based on these genes, most of the interactions between proteins were focused on ESR1, and functional pathways related to proteoglycans, immune cell signaling pathways, and apoptotic cell clearance were most enriched (Figure 7F-G).

## 4 Discussion

Breast cancer is a complex and diverse tumor type with a high degree of biological heterogeneity[7]. Accumulating evidence indicates that glucose metabolism had a critical role in tumor progression, controlling cell activities and proliferation and contributing to establishment of the TME, with eventual effects on patient survival [43]. Here, we integrated multi-omics analysis and *in vitro* experiments, evaluated the prognosis of breast cancer patients from glucose metabolism and TME perspectives, and comprehensively explored the role of PMAIP1 in the development of breast cancer.

The “Warburg effect” implicates abnormal glucose metabolism activity in cancer development, and infiltration of immune cells in the TME is another significant factor. Higher levels of infiltration are often associated with better survival. In the TCGA-BRCA dataset, we identified SORBS1, SIRT7, PMAIP1, and PGK1 as significant glucose-metabolism-related hub genes; these, together with selected protective immune cells, were used to construct a GLC-TME combined prognostic model. This integrated prognostic model showed optimized prediction ability and accuracy. When breast cancer patients were divided into four groups with significantly differential survival outcomes, the GLC-TME classifier was shown to function as an independent risk factor.

Functional enrichment analysis was also performed to identify the activated pathways in different GLC-TME subgroups. Vascular development and angiogenesis, as well as glycoprotein-related pathways, were specifically enriched in the group with the poorest survival (GLC-High+TME-Low). Furthermore, the group with the best survival (GLC-Low+TME-High) showed enrichment of pathways related to positive regulation of the immune system, including immune cell differentiation, immune memory process, and leukocyte adhesion. Pathways related to chemokine production, leukocyte adhesion, and propytosis were deactivated in the poor-survival group, consistent with the promotion of immune cell recruitment in GLC-Low+TME-High group, as determined by TIP analysis.

Microvessel development is closely associated with glucose metabolism in breast cancer, especially in the TME. The stroma in the TME contains fibroblasts, immune cells, and other components, and the glucose-metabolic interactions between cancer cells and stromal cells promote tumor growth, progression, angiogenesis, and metastasis [44, 45]. PFKFB3, a glycolysis-related enzyme, regulates vascular endothelial growth factor, thereby regulating the angiogenesis process in the TME [46]. In addition, aerobic glycolysis provides substrates for glycosylation in breast cancer, contributing to the establishment of the characteristic microenvironment of proliferation and metastasis of cancer cells [47].

Stimulated by antigens led to a marked increase in proliferation of T cells, consistent with upregulated glucose metabolism in these cells [48]. However, the excessive cellular activities in tumor cells consume large amounts of glucose in the environment, thus declined the proliferation process of T cells [49]. In addition, the glycolysis process produces lactate, which blocks lactate exportation of T cells and thus directly suppresses CD8+ T cell proliferation and cytotoxic activity [50]. CD4+ T memory cells are an important group of antigen-specific CD4+ T cells; they provide a persistent and specific immune response that is also dependent on glucose support and glycolysis [51, 52]. Lack of glucose may thus decrease the activity of CD4+ T memory cells and contribute to the development of cancer. Given the essential role of glucose metabolism in shaping the TME and regulating immune responses, targeting glucose-metabolism-related targets may offer promising therapeutic strategies for breast cancer treatment.

To locate critical glucose-metabolism-related targets, external dataset analysis and qRT-PCR experiments were conducted. The results showed differential expression levels of significant hub genes, especially PMAIP1. PMAIP1 encodes a BCL-2 protein family member that functions as a p53 downstream target in cells. Previous research reported in lung cancer and pancreatic cancer [53], the protein encoded by PMAIP1 has been implicated in glucose-deprivation-induced cell death and induces apoptosis after glucose depletion [54]. Under glucose limitation, PMAIP1 inhibits the anti-apoptotic function of the anti-apoptotic protein MCL-1 by competitively binding to it. The up-regulation of PMAIP1 weakened the dependence of tumor cells on anti-apoptotic proteins, such as MCL-1, and thus promoted apoptosis. This implies that PMAIP1 makes tumor cells more susceptible to apoptosis induction and more sensitive to glucose metabolism pathway inhibitors when glucose metabolism is dysregulated by regulating apoptosis-related signaling pathways.

In mitochondria, PMAIP1 prompts cytochrome c release and activates the caspase cascade reaction, leading to cleavage of multiple apoptosis-related substrates accompanied by loss of mitochondrial membrane potential [55]. Impairment of mitochondrial function also leads to increased generation of reactive oxygen species (ROS), further exacerbating oxidative stress [56]. In the endoplasmic reticulum (ER), PMAIP1 further exacerbates ER stress by interfering with calcium homeostasis [57]. Together, these stresses activate apoptosis signal-regulated kinase 1 (ASK1), which in turn initiates downstream MAPK signaling pathways, particularly p38 and JNK [58]. Activation of these kinases subsequently prompts the activation of the transcription factors AP-1 and ATF-2, which induce the expression of pro-apoptotic genes, thus amplifying the apoptotic effects induced by PMAIP1 [59].

Therefore, drugs that can promote PMAIP1 expression or prevent the degradation of its encoded protein may have potential anti-tumor effects by modulating glucose metabolism in breast cancer and provide new therapeutic targets for metabolic intervention and immunotherapy.

To further investigate the role of PMAIP1 in breast cancer, in vitro functional experiments in MCF-7 lines indicated that the knockdown of PMAIP1 led to the increased proliferation, migration and invasion of breast cancer. Meanwhile, the transcriptomic analysis further demonstrated an upregulation in glucose metabolism after PMAIP1 knockdown. Compared to NC group, PMAIP1-KD group showed upregulated canonical glucose metabolism related genes, such as G6PC3, PKFP, PPARA and PIK3CA, indicating the variated metabolism status [60–63]. The differential overall glucose metabolism landscape in two groups suggested that PMAIP1 may play a critical role in regulating both tumor growth and glucose metabolism in breast cancer.

Furthermore, single-cell analysis was used to explore glucose metabolism in breast cancer. The distribution of PMAIP1 was mainly concentrated in luminal cells and immune cells, including T cells and macrophages, which might suggest the potential regulation of PMAIP1 both in tumor cells and immune cells in breast cancer. Luminal cells were identified as the pathogenic cell clusters based on integrated single-cell data and breast cancer GWAS data, indicating that luminal cells might have a critical role in the development of breast cancer. Thus, luminal cells were specifically extracted and re-clustered; then, the “AddModuleScore” function was used to evaluate the expression of significant genes, as well as genes related to cancer stemness. Cluster 0 was assigned as the root node of the pseudotime, as the specific cluster showed the highest cell stemness. Clusters 1 and 4 showed increased proportions in the Score-high group, and pseudotime analysis revealed that they belonged to the same heterogenetic differentiation branch of luminal cells. Located in the terminal differentiation stage, cluster 1 showed increased functions related to energy metabolism, cell respiratory and redox reactions, which are all glucose-metabolism-related pathways. Generally, cluster 1 represents pathogenic cells in breast cancer that are effectively regulated by significant genes, eventually resulting in upregulation of glucose metabolism processes. Therapies targeting these genes could be used to regulate the differentiation of glucose-metabolism-related luminal cells; this could make them effective means of inhibiting or even reversing the progression of abnormal glucose metabolism in breast cancer.

It is crucial to understand the interactions between PMAIP1+ luminal cells and the immune microenvironment. Intercellular communication analysis revealed interactions between PMAIP1+ luminal cells and macrophages through the MIF and galectin pathways. MIF activates CD74, a signaling receptor that triggers the activation of ERK/MAPK, JAB1/AP-1, and NF-κB, eventually promoting the M1 polarization and infiltration of macrophages in the TME [64, 65]. M1-polarized macrophages secrete TNF-α, which directly targets cancer cells, thereby promoting inflammatory responses, killing intracellular pathogens, and contributing to immune resistance to tumor development [66]. Furthermore, galectin-12 and intracellular galectin-3 positively regulate macrophage phagocytosis [67]. Extracellular vesicles derived from tumor cells and immune cells mediate mutual communication at proximal and distal sites, and extracellular vesicles in the TME show immunostimulatory molecular profiles associated with a Th1/M1 polarization signature, stimulating antitumor immunity [68]. Here, the enhanced intensity of MIF and galectin communications observed between PMAIP1+ luminal cells and macrophages indicate promotion of the immune response of M1 macrophages in breast cancer, resulting in increased immune infiltration and tumor killing effects, consistent with the protective role of M1 macrophages identified in the immune infiltration and multivariate Cox analyses.

The metabolic pathway analysis indicated upregulated glycosylation and glycosphingolipid biosynthesis in PMAIP1− luminal cells, compared to PMAIP1+ luminal cells. In breast cancer, dysregulated glycosylation is associated with decreased treatment effects and increased tumor malignancy; for example, the glycosylation status of PD-L1 affects response to immune checkpoint blockade therapy in triple-negative breast cancer [69]. In addition, the immune response of B cells is inhibited by glycophospholipid structures on the surface of tumor cells [70], consistent with the decreased activity of the majority of metabolic pathways in B cells.

Furthermore, the protective effects of PMAIP1 were explored by Bayes deconvolution. Patients with higher proportions of PMAIP1+ luminal cells had better survival outcomes, and the varying proportions of PMAIP1+ luminal cells among different clinical subtypes suggest the prognostic value of PMAIP1+ luminal cells in clinical practice.

The proportion of PMAIP1+ luminal cells could also be related to tumor phenotypes. For instance, patients in the low PMAIP1+ luminal cell group may have more TP53 mutations. TP53 regulates the cell cycle, apoptosis, DNA repair, and tumor suppression [71], and its increased mutation is associated with higher tumor malignancy, drug resistance, and poorer prognosis. In addition, the immune analysis revealed better T cell infiltration, T cell function, and immune therapy response in the high PMAIP1+ luminal cell group, and the results of the drug sensitivity analysis indicated that higher proportions of PMAIP1+ luminal cells were correlated with greater sensitivity to drugs including cisplatin, axitinib, and gefitinib. The high drug sensitivity of the high PMAIP1+ luminal cell group suggests that the regulation of glucose metabolism by PMAIP1 in breast cancer affects diverse biological processes, including DNA repair, vascular development, and the epidermal growth factor receptor (EGFR) signaling pathway, further demonstrated the potential of PMAIP1 as a clinical treatment target.

The WGCNA method was used to identify the co-expressed gene modules in breast cancer that were most related to PMAIP1+ luminal cells. ESR1, which is the receptor of estrogen in breast cancer, showed the most frequent interactions with other proteins. ESR1 inhibits the function of EGFR and human EGFR2 (HER2) receptor tyrosine kinase, thereby inhibiting the epithelial–mesenchymal transition of cells and inhibiting the distant metastasis of breast cancer [72, 73]. PMAIP1 could promote this inhibitory effect, leading to better prognosis.

Functional enrichment of the intersected genes showed that they were enriched in pathways related to immune cell signaling and apoptotic cell clearance. Based on the results of the immune therapy prediction conducted with TIDE, we speculate that PMAIP1 promotes infiltration and response of immune cells through regulating the glucose metabolism of cancer cells. In particular, Fc-γR-mediated phagocytosis was enhanced in PMAIP1+ luminal cells, consistent with the positive interaction between PMAIP1+ luminal cells and macrophages in breast cancer [74]. In addition, PMAIP1 is a member of the BCL-2 family, which has strong association with cell apoptosis [75]. The activation of the apoptotic cell clearance process indicated increased intensity of cell apoptosis and phagocytosis, further emphasizing the critical role of PMAIP1 in regulating cancer cell apoptosis and promoting a positive immune response in breast cancer.

However, there were some limitations. Although the GLC-TME score was found to be an independent risk factor in breast cancer, different subtypes of breast cancer were not considered. Accommodating individual variance among patients is important in personalized medicine. Moreover, although we used in silico analysis and in vitro experiments to comprehensively explore the functions of PMAIP1 in breast cancer, more in-depth mechanistic research is needed to thoroughly elucidate its precise roles and significance in the prognostic landscape of breast cancer patients.

## 5 Conclusion

In summary, in the present study, we identified four significant prognostic genes related to glucose metabolism in breast cancer, established a combined prognostic model, and further explored the function of PMAIP1 in breast cancer. Through bulk-seq, transcriptomic analysis, and in vitro experiments, PMAIP1 was confirmed to be a suppressor of tumor proliferation, invasion, migration, potentially by regulating glucose metabolism. Moreover, single-cell analysis and Bayes deconvolution revealed a specific role of PMAIP1+ luminal cells in breast cancer, with a higher proportion of PMAIP1+ luminal cells indicating better survival, immune function, and curative effect of drugs.

## 7 Author Contributions

YZ, QY and LZ designed the research. LZ and HX supervised the research. YZ collected data. QY and YZ performed data analysis; XH conducted the experiment validation. YZ drafted the manuscript. LZ, HX and QY revised the manuscript. All authors have given approval to the publication of the article.

## 9 Abbreviations

ATP: Adenosine Triphosphate
TCA: Triple Citric Acid
OXPHOS: Oxidative Phosphorylation
TME: Tumor Micro-environment
TCGA-BRCA: The Cancer Genome Atlas Breast Invasive Carcinoma
TPM: Transcripts Per Million Reads
METABRIC: Molecular Taxonomy of Breast Cancer International Consortium
GEO: Gene Expression Omnibus
GSEA: Gene Set Enrichment Analysis
DEG: Differentiated Expressed Gene
LASSO: Least Absolute Shrinkage and Selection Operator
SD: Standard Deviation
ROC: Receiver Operating Characteristic
WGCNA: Weighted Correlation Network Analysis
NES: Normalized Enrichment Score
TIP: Tracking Tumor Immunophenotype
qRT-PCR: quantitative real-time polymerase chain reaction
PCA: Principal Component Analysis
UMAP: Uniform Manifold Approximation and Projection
L-R: Ligand-Receptor
KEGG: Kyoto Encyclopedia of Genes and Genomes
IC50: half-maximal inhibitory concentration
TGF-β: Transforming growth factor β
VEGF: vascular endothelial growth factor
TRS: trait-relevant score
L-R: ligand-receptor
EM: Expectation-maximization
SNV: simple nucleotide variation
TIDE: Tumor Immune Dysfunction and Exclusion
IC50: half-maximal inhibitory concentration
GDSC: Genomics of Drug Sensitivity in Cancer
PPI: protein-protein interaction
HER2: human epidermal growth factor receptor 2
EMT: epithelial-mesenchymal transition.

## 10 Acknowledgments

Thanks to the patients who supplied clinical and sequencing data for this study.

## 11 Supplementary Material

**Figure S1.**
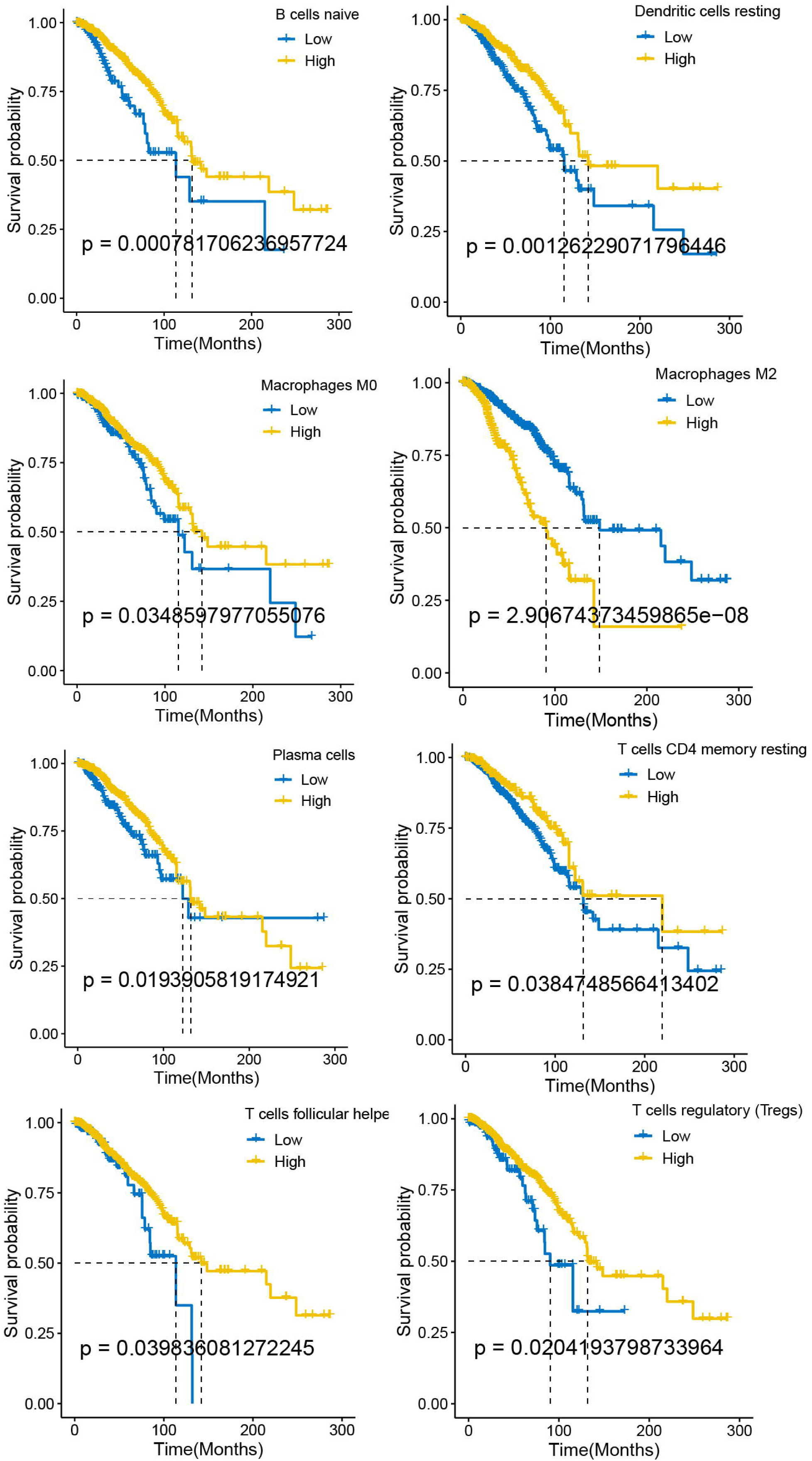
The Kaplan-Meier survival curves of patients with breast cancer from TCGA, grouped by immune cell proportion acquired from CIBERSORT. The threshold was set as p<=0.05;

**Figure S2.**
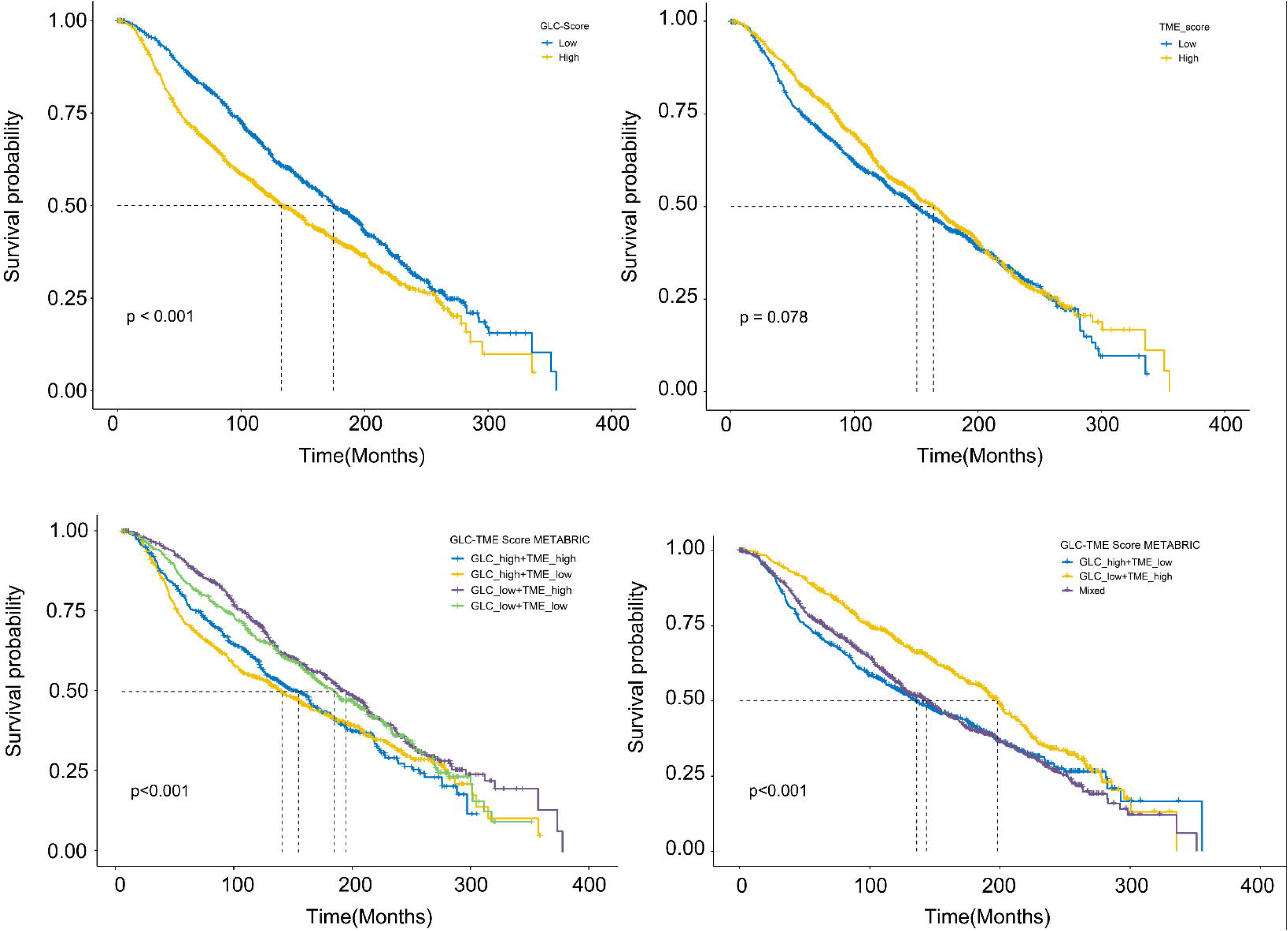
The Kaplan-Meier survival curves of patients with breast cancer from METABRIC cohort, for GLC-Score, TME-Score and GLC-TME prognostic model validation;

**Figure S3.**
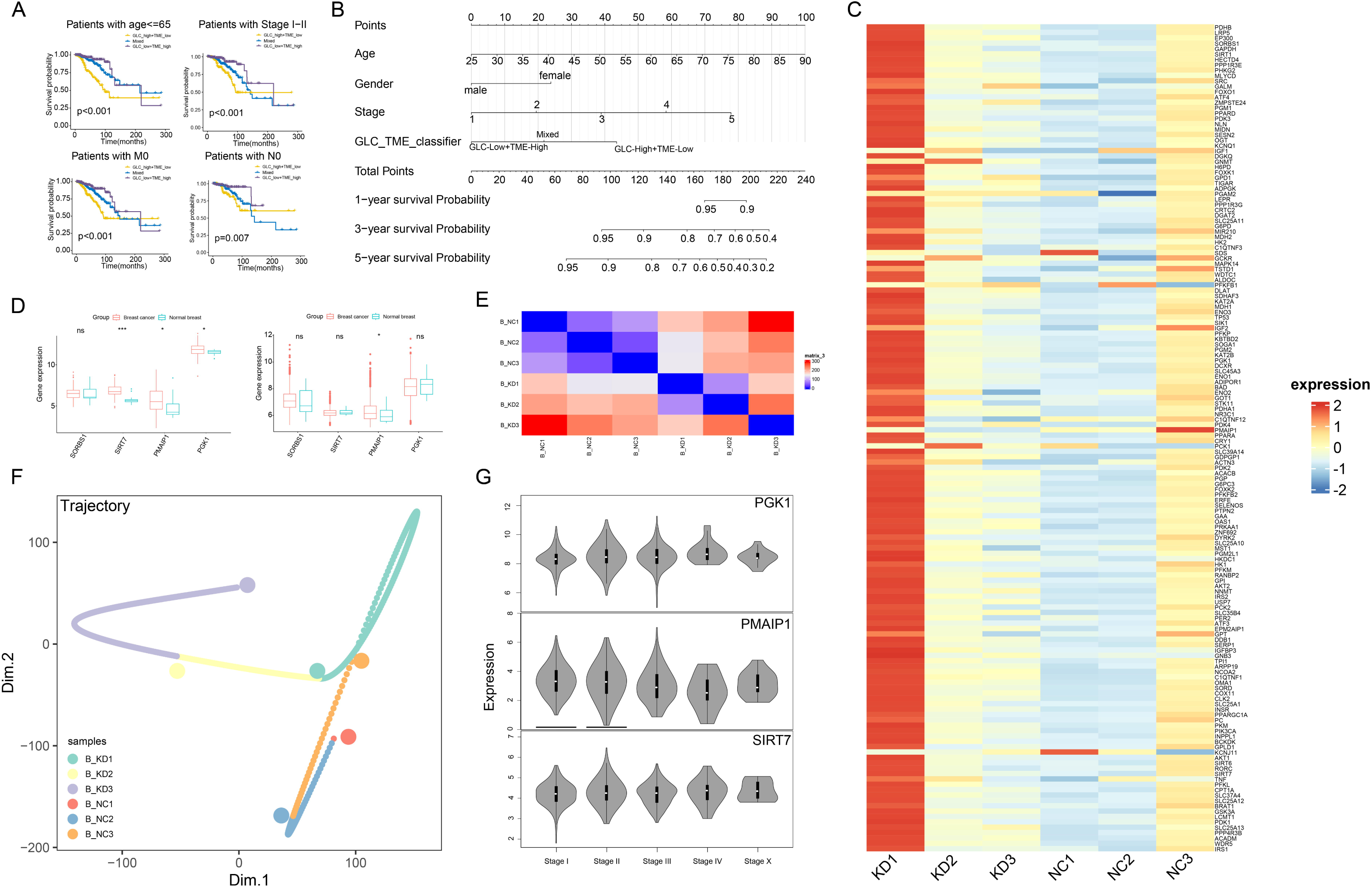
Survival analysis in breast cancer subtypes and external dataset pre-processing. (A) The survival differences of three groups in specific subtypes of breast cancer patients. (B) The nomogram established based on the combined prognostic model. (C) The expression variation of glucose metabolism related genes in MCF-7 and PMAIP1-KD MCF-7 groups (D) External validation of expression levels of 4 significant prognostic genes in METABRIC and GSE42568. (E) PCA distance between samples for bulk pseudotime analysis. (F) Differentiation trajectory of 6 samples. (G) Expression levels of 3 significant prognostic genes in different stages of breast cancer, acquired from GEPIA database.

**Figure S4.**
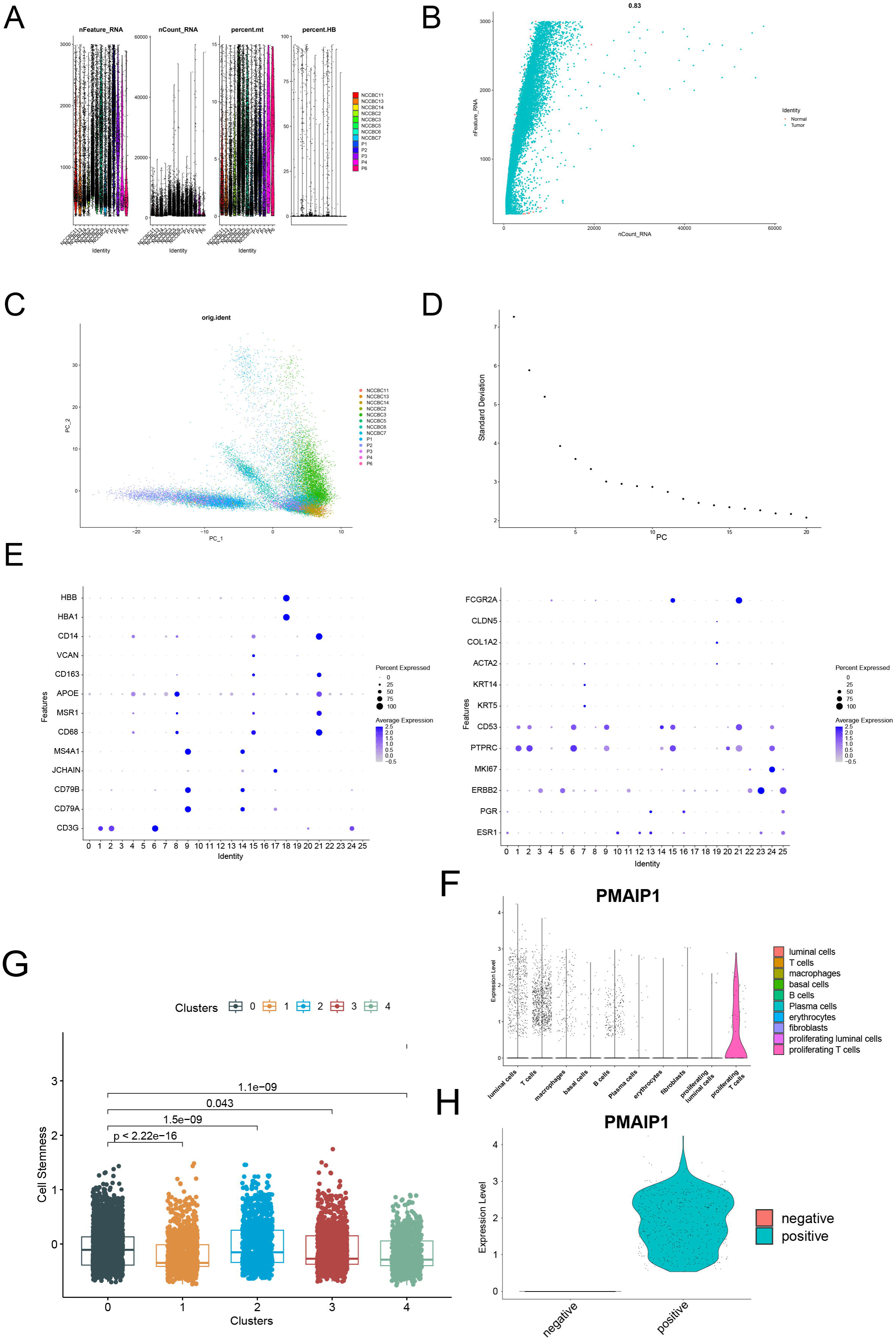
The quality control and data processing of single-cell data GSE195861. (A) Quality control of scRNA data samples. (B) Correlation analysis between nCount RNA and nFeature RNA. (C) Scatter plot showing the scores of individual cells (points) along the top two principal components. (D) Standard deviation (y-axis) accounts for top 20 PCs (x-axis) to identify the number of significant PCs based on the presence of an “elbow”. Approximately 10 PCS are chosen for the analysis. (E) Dotplot showing the average expression level of canonical marker genes of each cluster in scRNA samples. (F) PMAIP1 expression in different cell clusters. (G) Differential cell stemness among luminal clusters, calculated by “AddModuleScore” function. (H) PMAIP1 expression in PMAIP1+ luminal cells and PMAIP1-luminal cells.

**Figure S5.**
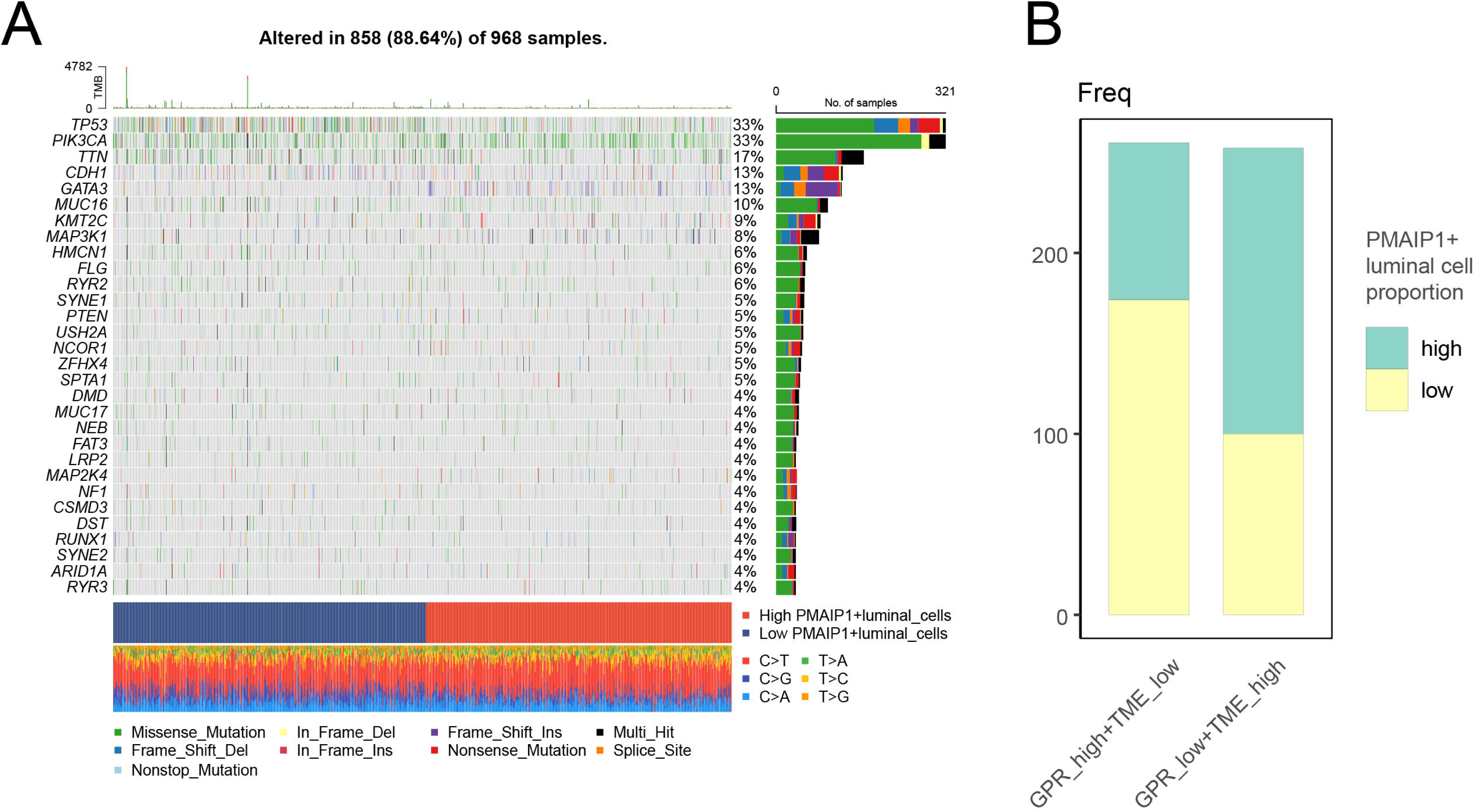
Mutation and patient proportion of PMAIP1+ luminal cell groups. (A)The mutation landscape of breast cancer patients, grouped by the PMAIP1+ luminal cell proportion. (B) The proportion of high/low PMAIP1+ luminal cell patients in prognosis-related GLC-TME groups.

**Figure S6.**
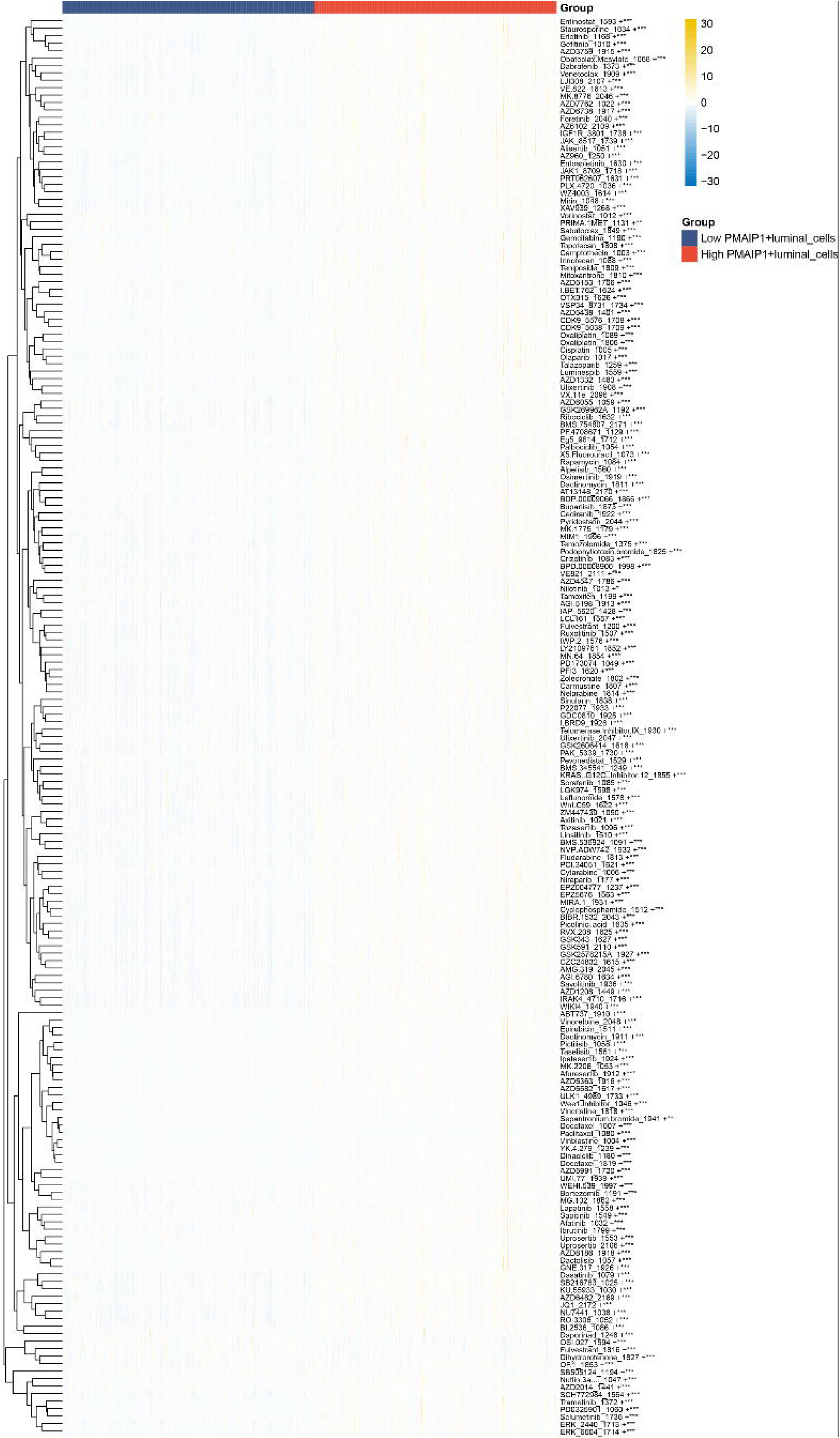
Overall landscape of drug sensitivity of patients in high and low PMAIP1+ luminal cell groups.

## 12 Data Availability Statement

The public datasets selected in our research can be found and downloaded for free online, while the databases we searched and accession number(s) are all provided in article or supplement materials. MCF-7 cell line mRNA sequenced data will be available upon reasonable request to Yidong Zhang.

