## Supplement Material for "PMAIP1-Mediated Glucose Metabolism and its Impact on the Tumor Microenvironment in Breast Cancer: Integration of Multi-Omics Analysis and Experimental Validation"

Supplementary Material

### 1 Supplementary Figures and Tables

#### 1.1 Supplementary Figures

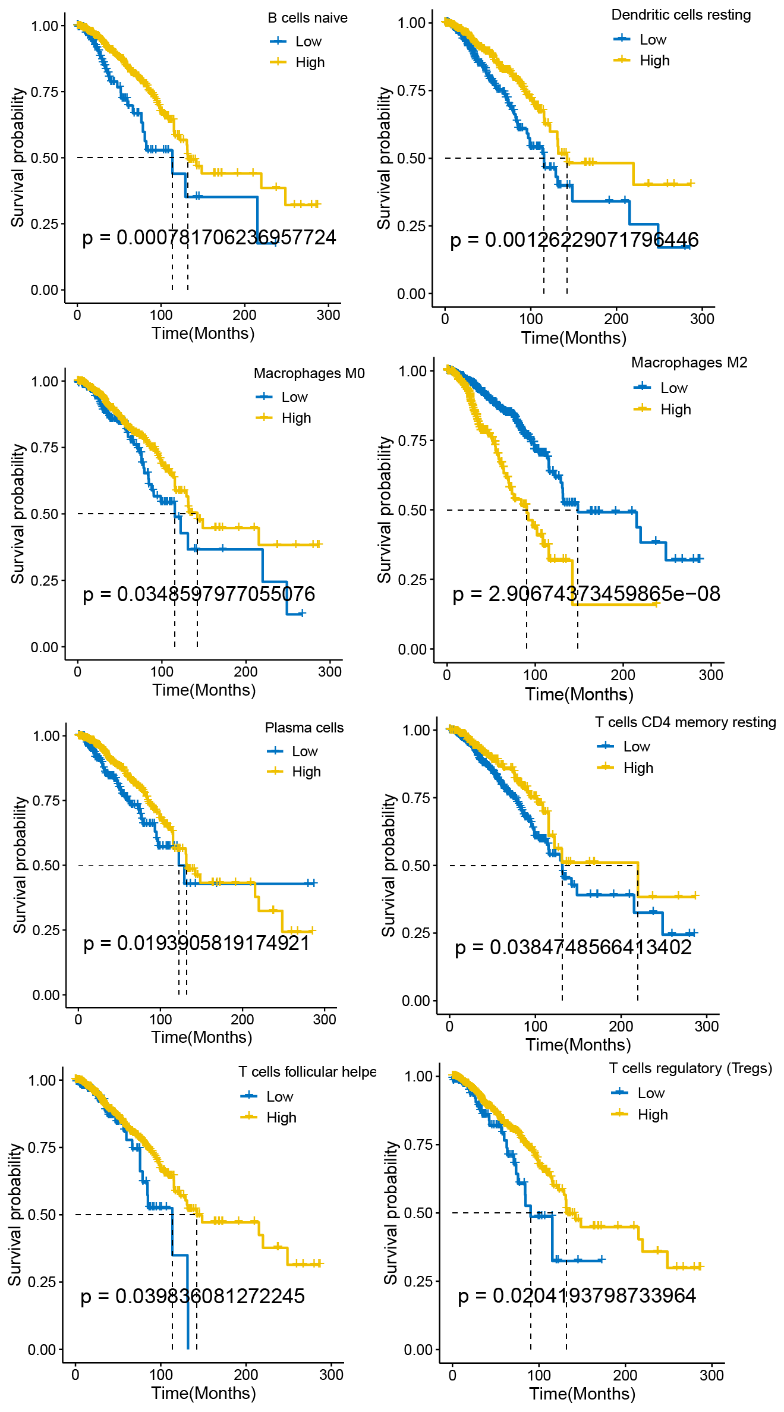

**Figure S1** The Kaplan-Meier survival curves of patients with breast cancer from TCGA, grouped by immune cell proportion acquired from CIBERSORT. The threshold was set as p<=0.05.

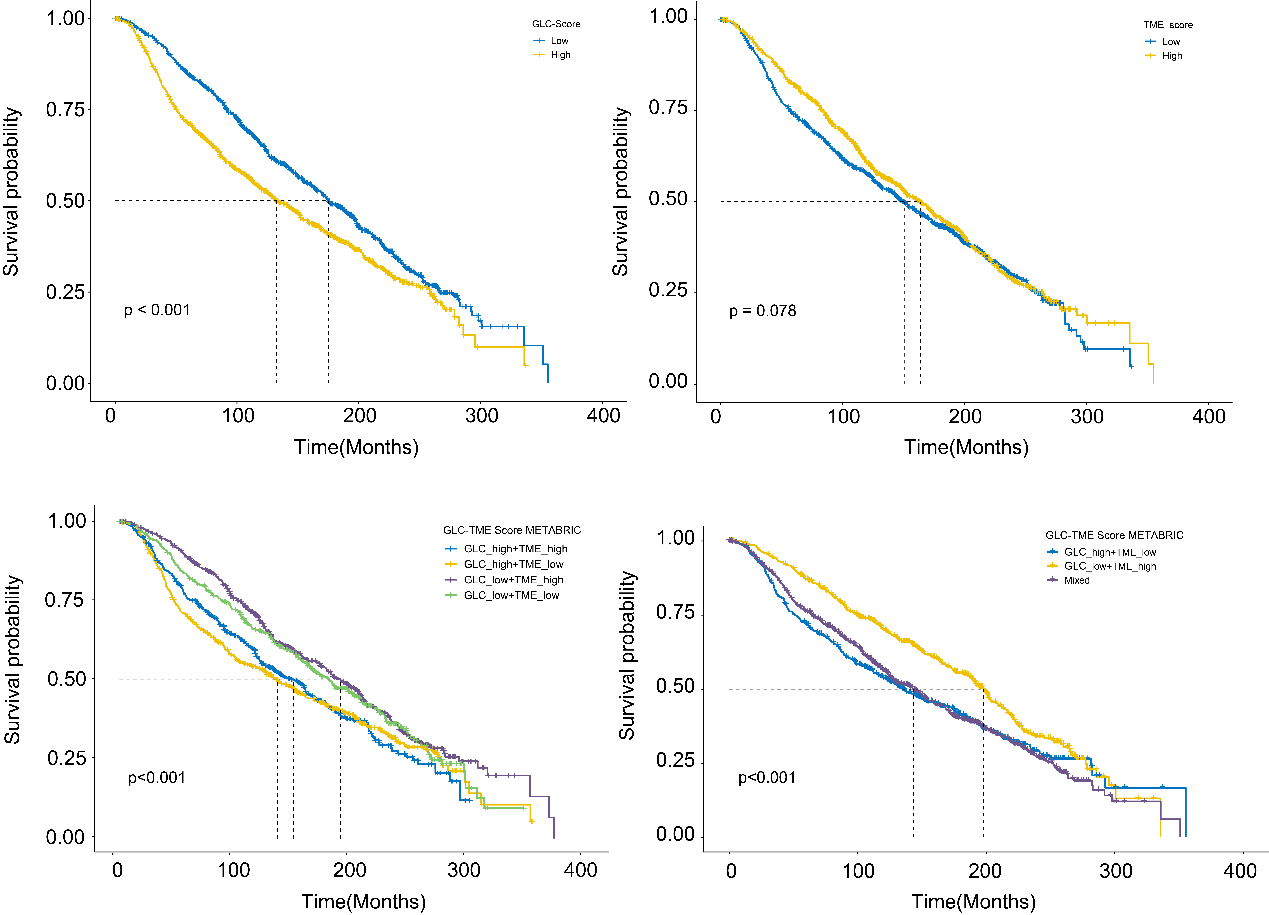

**Figure S2** The Kaplan-Meier survival curves of patients with breast cancer from METABRIC cohort, for GLC-Score, TME-Score and GLC-TME prognostic model validation.

**Figure S3** Survival analysis in breast cancer subtypes and external dataset pre-processing. (A) The survival differences of three groups in specific subtypes of breast cancer patients. (B) The nomogram established based on the combined prognostic model. (C) The expression variation of glucose metabolism related genes in MCF-7 and PMAIP1-KD MCF-7 groups. (D) External validation of expression levels of 4 significant prognostic genes in METABRIC and GSE42568. (E) PCA distance between samples for bulk pseudotime analysis. (F) Differentiation trajectory of 6 samples. (G) Expression levels of 3 significant prognostic genes in different stages of breast cancer, acquired from GEPIA database.

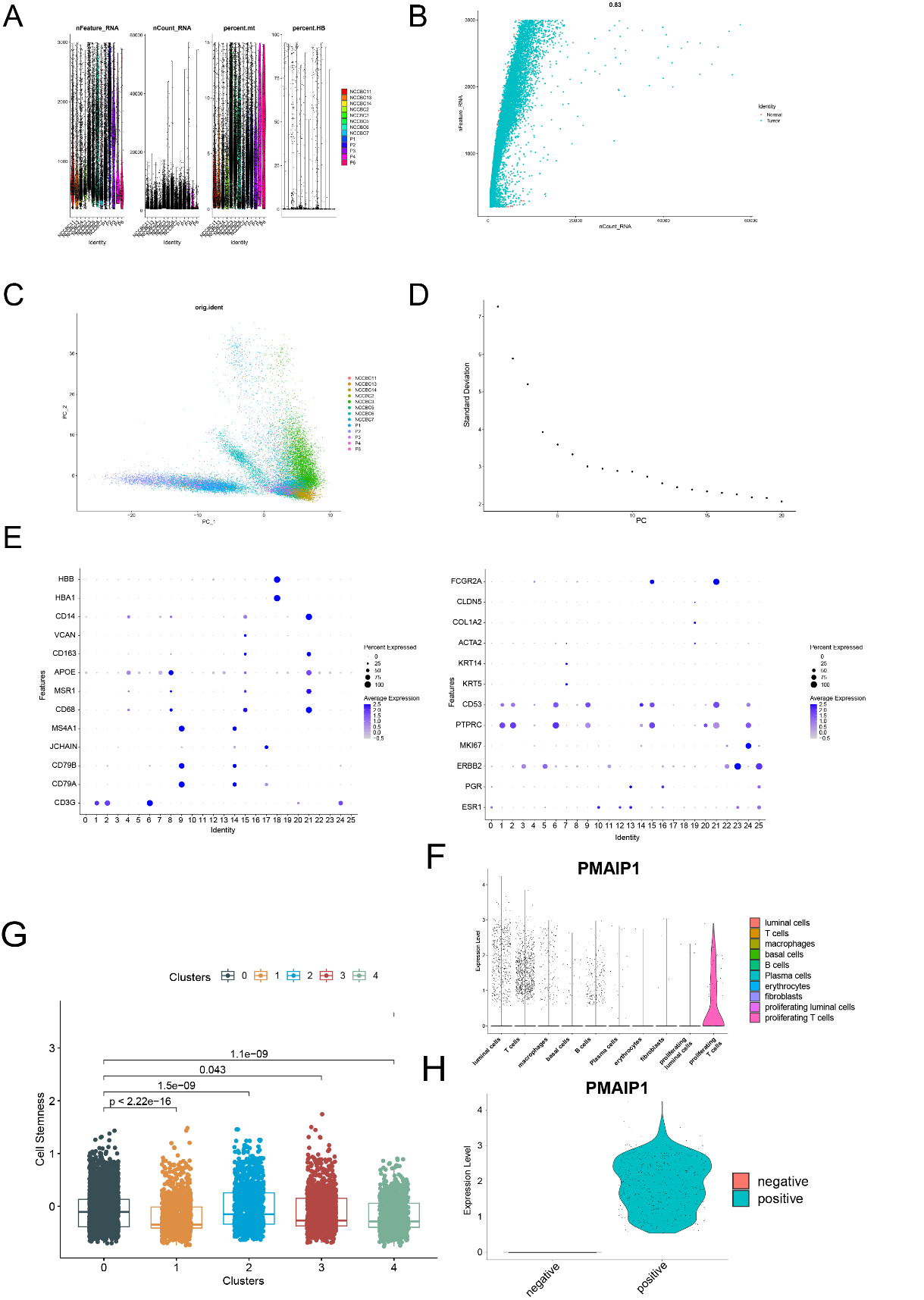

**Figure S4** The quality control and data processing of single-cell data GSE195861. (A) Quality control of scRNA data samples. (B) Correlation analysis between nCount RNA and nFeature RNA. (C) Scatter plot showing the scores of individual cells (points) along the top two principal components. (D) Standard deviation (y-axis) accounts for top 20 PCs (x-axis) to identify the number of significant PCs based on the presence of an “elbow”. Approximately 10 PCS are chosen for the analysis. (E) Dotplot showing the average expression level of canonical marker genes of each cluster in scRNA samples. (F) PMAIP1 expression in different cell clusters. (G) Differential cell stemness among luminal clusters, calculated by “AddModuleScore” function. (H) PMAIP1 expression in PMAIP1+ luminal cells and PMAIP1- luminal cells.

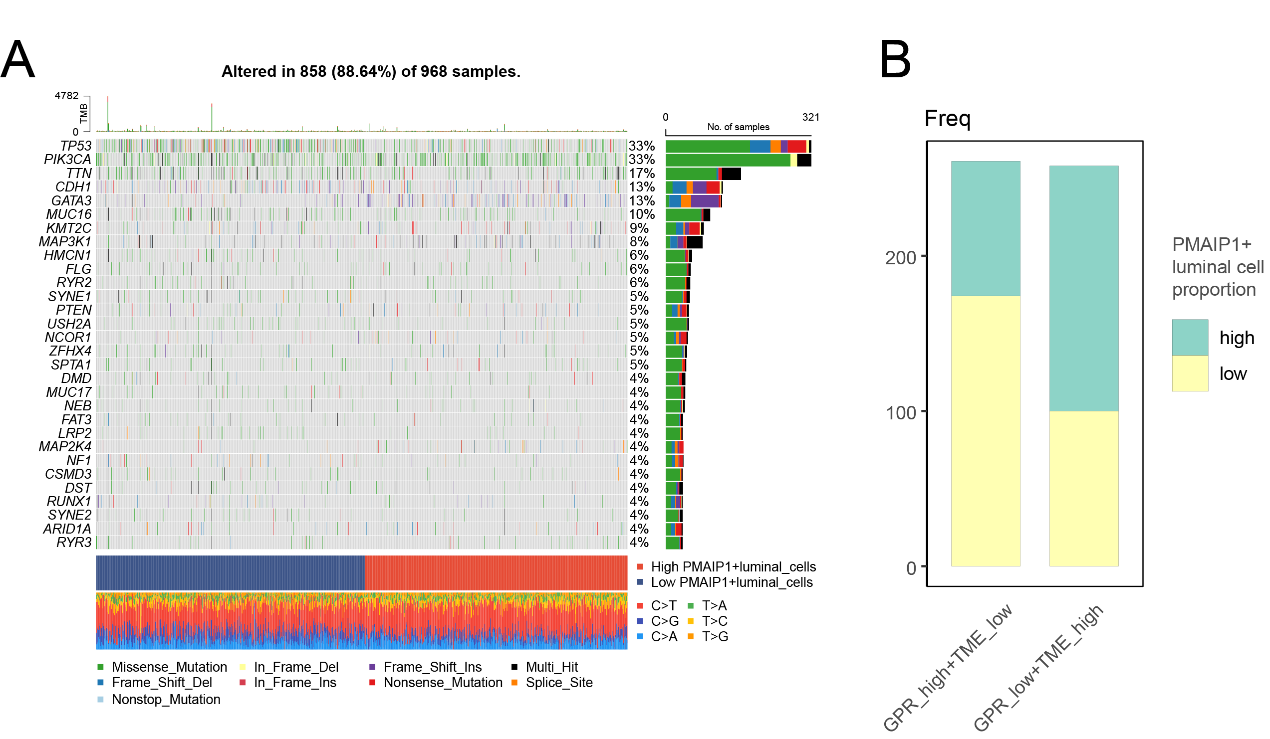

**Figure S5** Mutation and patient proportion of PMAIP1+ luminal cell groups. (A)The mutation landscape of breast cancer patients, grouped by the PMAIP1+ luminal cell proportion. (B) The proportion of high/low PMAIP1+ luminal cell patients in prognosis-related GLC-TME groups.
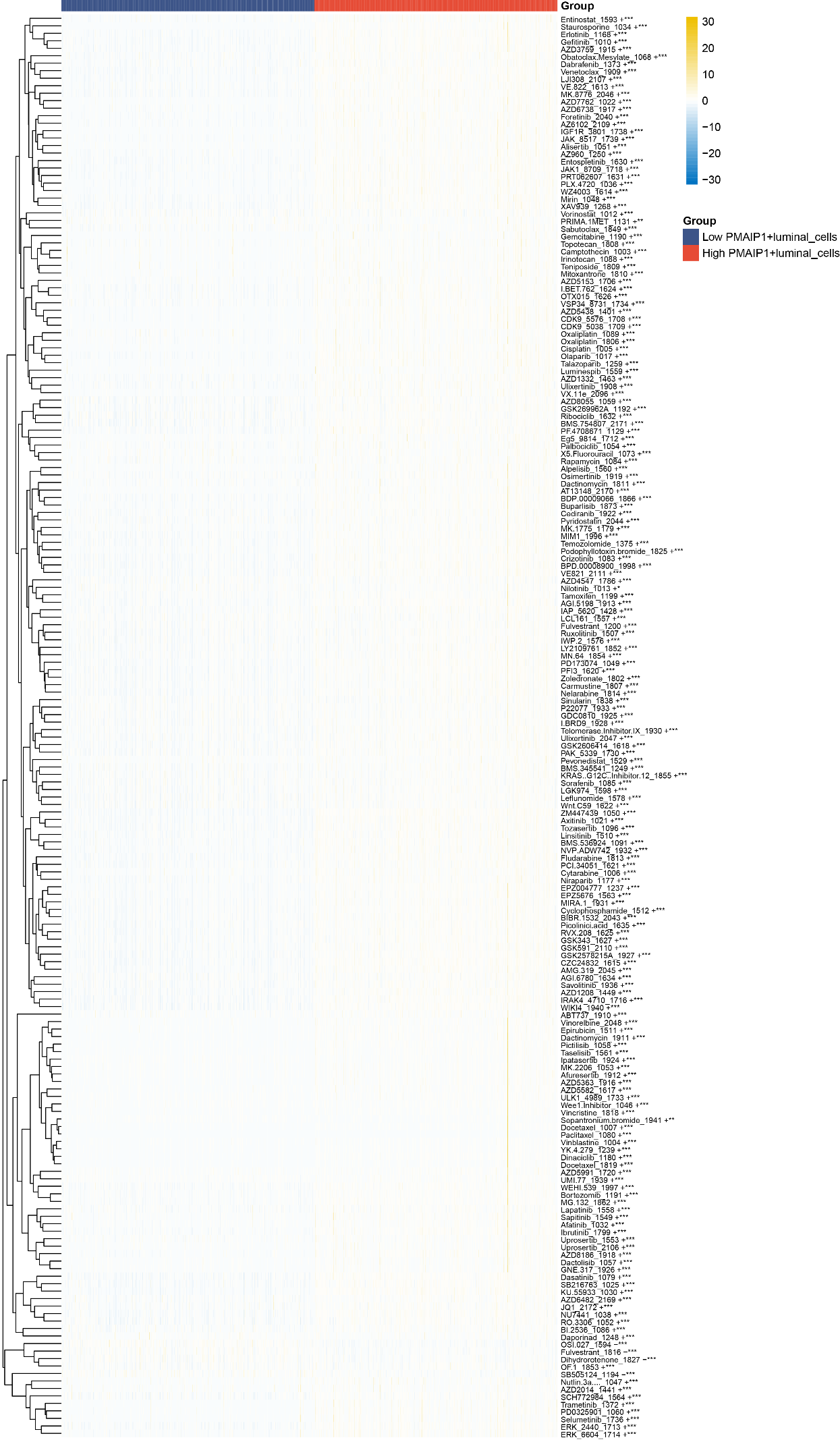

**Figure S6** Overall landscape of drug sensitivity of patients in high and low PMAIP1+ luminal cell groups.

#### 1.2 Supplementary Table

**Supplementary Table1**. Sequences of primers used for qRT-PCR in this study

| Primer | Sequence |
| --- | --- |
| SIRT7 (F) | 5'- CAGGGAGTACGTGCGGGTGT -3'  5'- TCGGTCGCCGCTTCCCAGTT -3' |
| SIRT7 (R) |  |
| PGK1 (F) | 5'- TGGAGTCGAGCACTCACAAC -3' |
| PGK1 (R) | 5'- GTATAGGCCATTCCTCCGCC -3' |
| PMAIP1 (F) | 5'- GAGCAGAAGAGTTTGGATATCAGA -3' |
| PMAIP1 (R) | 5'- GCAAGAACGCTCAACCGA -3' |
| GAPDH (F) | 5'- GTCTCCTCTGACTTCAACAGCG -3' |
| GAPDH (R) | 5'- ACCACCCTGTTGCTGTAGCCAA -3' |

**Supplementary Table2**. Target sequences of siRNAs for each gene

| siRNA targeting PMAIP1 |  | Sequence |
| --- | --- | --- |
| hs-PMAIP1-si-1 | sense (5'-3') | AGUCGAGUGUGCUACUCAAdTdT  UUGAGUAGCACACUCGACUdTdT |
| hs-PMAIP1-si-1 | antisense(5'-3') |  |
| hs-PMAIP1-si-2 | sense (5'-3') | GGCGCGCAAGAACGCUCAAdTdT |
| hs-PMAIP1-si-2 | antisense(5'-3') | UUGAGCGUUCUUGCGCGCCdTdT |
| hs-PMAIP1-si-3 | sense (5'-3') | GAAACUUCUGAAUCUGAUAdTdT |
| hs-PMAIP1-si-3 | antisense(5'-3') | UAUCAGAUUCAGAAGUUUCdTdT |
